# Engineering Extracellular Vesicle Production through Magnetic Ion Channel Activation for Bone Regeneration

**DOI:** 10.1101/2025.08.07.669024

**Authors:** Afeesh Rajan Unnithan, Kenny Man, Kritika, Lee A. Gethings, Christopher J. Hughes, Alicia Keenan, Liam Heaney, Sophie C. Cox, Owen G. Davies, Alicia J. El Haj

**Affiliations:** Faculty of Life Sciences, Centre for Pharmaceutical Engineering Science, School of Pharmacy and Medical Sciences, University of Bradford, Bradford, UK; Department of Oral and Maxillofacial Surgery & Special Dental Care, University Medical Center Utrecht, Utrecht, 3508 GA The Netherlands; Regenerative Medicine Center Utrecht, University Medical Center Utrecht, Utrecht, The Netherlands; Department of Chemistry, University of Delhi, Delhi-110007, India; Waters Corporation, Wilmslow; School of Sport, Exercise and Health Sciences, Loughborough University; School of Chemical Engineering, University of Birmingham, Birmingham, B15 2TT UK; Healthcare Technologies Institute, Institute of Translational Medicine, School of Chemical Engineering, University of Birmingham, Birmingham, United Kingdom

**Keywords:** extracellular vesicles, magnetic nanoparticles, nanomedicine, mechanotransduction, osteogenesis, bioengineering

## Abstract

Bone disorders represent a significant global health challenge. Extracellular vesicles (EVs) are emerging as a promising nanotherapeutic approach for bone regeneration, addressing the translation barriers associated with cell-based therapies. Despite their immense potential, the clinical application of EVs is limited by low production yields and inconsistent quality. Magnetic Ion Channel Activation (MICA) utilises remote magnetic fields to stimulate mechano-sensitive ion channels through magnetic nanoparticles (MNPs). This study explores the potential of utilising MICA to enhance the production yield and therapeutic efficacy of EVs for bone regeneration. The findings demonstrate that MICA significantly increased the production yield of EVs from MC3T3 pre-osteoblasts compared to magnetic stimulation or TREK1 functionalised graphene oxide-MNP particles alone. The obtained EVs exhibited typical size distribution, morphology, and EV protein expression consistent with nano-sized vesicles. Furthermore, MICA/TREK EVs treatment considerably enhanced human bone marrow-derived mesenchymal stem cells osteogenic differentiation and mineralisation compared to EVs derived from MICA, TREK, or untreated groups. Proteomics analysis revealed the enrichment of proteins involved in mechanotransduction and osteogenic differentiation within MICA/TREK EVs. In summary, these findings highlight the substantial potential of MICA as a platform to enhance the scalable production and therapeutic application of pro-regenerative EVs for bone augmentation strategies.

## 1. Introduction

Bone-related disorders, including traumatic injuries, osteoporosis and tumour resection defects play a significant clinical and socioeconomic burden globally. Osteoporosis is a highly prevalent disease and results in massive costs both to the individual and to society through associated fragility fractures. An estimated 10 million people over the age of 50 years have osteoporosis, and around 1.5 million fragility fractures occur in these patients each year[1]. Osteoporosis treatments aim to strengthen bones and reduce fracture risk, often involving medications like bisphosphonates, denosumab, or anabolic agents, alongside lifestyle changes like exercise and a calcium and vitamin D-rich diet. Current treatments often fail to fully restore bone mass and function and are focused on slowing down bone loss rather than stimulating new bone formation in critical areas such as the vertebrae. This highlights the urgent need for advanced therapeutic strategies that address restoring bone biological function following regeneration[2–4].

Increasing evidence has demonstrated the importance of cell-derived bioactive molecules in facilitating cellular communication and modulating diverse biological processes[5–7]. Among these bioactive factors, extracellular vesicles (EVs) have emerged as vital mediators of intracellular communication, playing pivotal roles in a variety of physiological and pathological processes including bone regeneration and immune modulation[8, 9]. EVs are cell-secreted lipid nanoparticles enriched with bioactive molecules such as proteins, nucleic acids and metabolites making them promising candidates for nanotherapeutic applications, especially in regenerative medicine[10, 11]. EV-based therapeutics offer significant advantages to traditional cell-based therapies including reduced immunogenicity, improved stability and the ability to cross biological membranes (i.e. blood-brain barrier)[12]. However, despite their potential, the translation of EV-based therapies to the clinical arena is hindered due to their inherent therapeutic potency, low and variable production yield and unreliable manufacturing methods [13, 14]. Conventional EV production strategies including genetic modifications, application of external forces and the use of chemical reagents often suffer from limitations such as scalability issues, cellular stress and even compromised EV bioactivity[15, 16]. Hence, there is a significant unmet need to refine the culture conditions to enhance EV therapeutic potency and yield for bone augmentation strategies.

Recent developments in mechanobiology and bioengineering have given new opportunities for EV production, especially with ion channel activation emerging as a potential target for modulating cellular behaviour[17] and EV production [18]. Ion channels play a crucial role in maintaining cellular haemostasis by mediating the ion flux across the cell membranes and hence regulating the intracellular signalling pathways[19]. Mechano-sensitive ion channels are responsive to external mechanical stimuli providing a unique opportunity to modulate cellular activity and EV production [20–22]. However, approaches using chemical inducers or random mechanical forces can often result in inconsistent cell stimulation and inducing cell stress, ultimately detrimentally impacting the production of therapeutic EVs[23, 24]. Therefore, a targeted and reproducible method for ion channel activation is critical for improving EV yield and its bioactivity[25–27].

Magnetic ion channel activation (MICA) represents an innovative approach to improve EV production. Magnetic nanoparticles (MNPs) under remote magnetic fields offer a non-invasive and controllable means of stimulating mechano-sensitive ion channels to trigger cellular processes from outside the body [26, 28]. MICA is well aligned with scalable platforms such as bioreactor technologies used for EV production. Previously, we reported that TREK1 functionalised Graphene oxide-MNPs (TGMNPs) resulted in enhanced MSc differentiation into bone cells with enhanced calcification and bone matrix production in vitro. In addition, MICA-induced enhancement of bone growth has been demonstrated *in vivo* in rodent and sheep models[29]. MICA can be fine-tuned to deliver consistent stimuli across large cell populations, addressing one of the primary limitations of current EV production techniques. Moreover, this non-invasive approach also facilitates real-time monitoring and control ensuring high yields of functional EVs suitable for clinical applications.

This study investigates the potential of harnessing MICA as a novel strategy to enhance the production and therapeutic potency of EVs for bone regeneration. TGMNPs in conjunction with MICA were used to culture pre-osteoblasts and EVs were obtained from the conditioned medium and then characterised. EVs from MICA/TREK stimulated osteoblasts were administered to human bone marrow-derived mesenchymal stem cells (hBMSCs) to evaluate their osteogenic potency (Figure 1). Moreover, proteomics analysis was conducted to elucidate the mechanisms by which the MICA EVs impart their pro-osteogenic function. By bridging the gap between mechanobiology and regenerative medicine, this approach has the potential to revolutionise the EV manufacturing process and pave the way for next-generation EV-based therapies.

**Figure 1.**
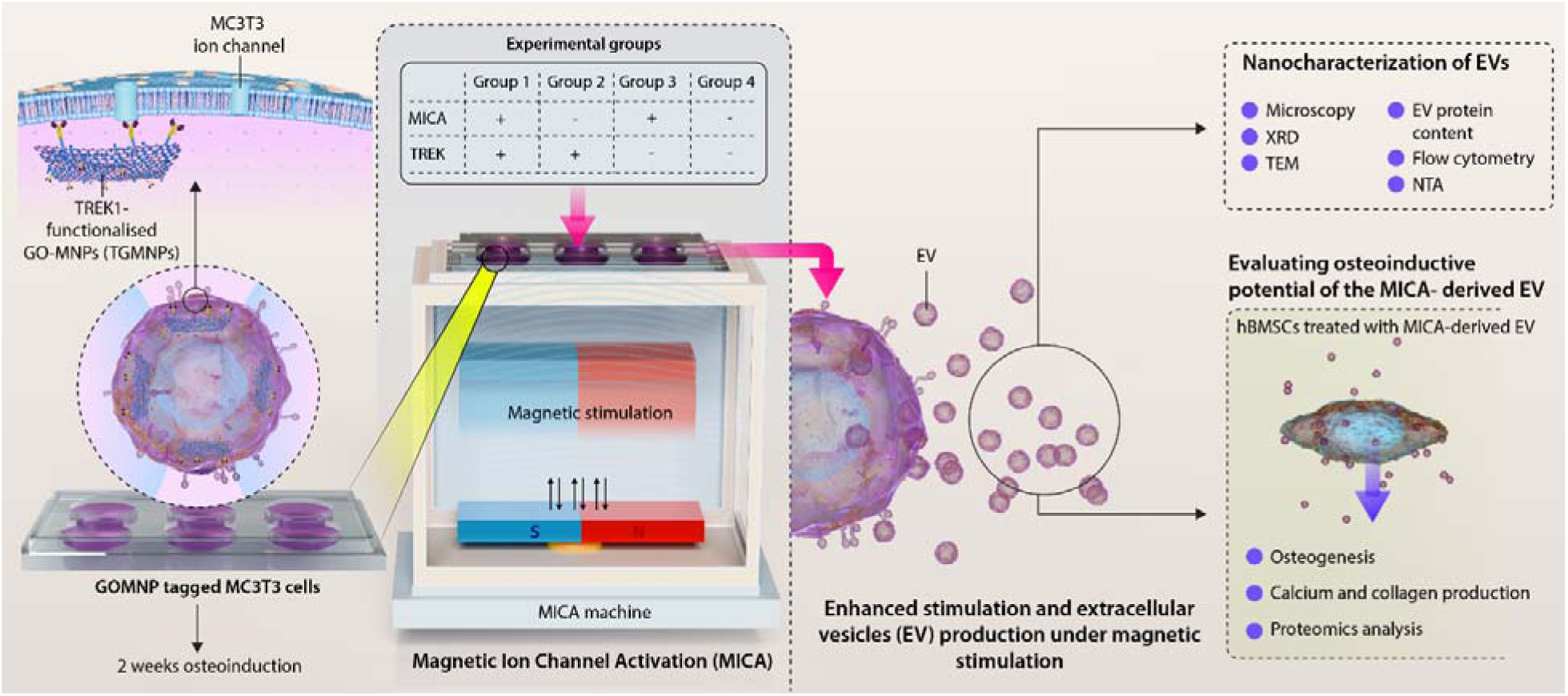
Experimental overview.

## 2. Materials and Methods

### Cell culture and reagents

MC3T3 pre-osteoblasts and hBMSCs were obtained from American Type Culture Collection (ATCC, UK) and Lonza (Lonza, UK) respectively. Basal culture media consisted of minimal essential medium (α-MEM; Sigma-Aldrich, UK) supplemented with 10% foetal bovine serum (FBS), 1% penicillin/streptomycin (Sigma-Aldrich, UK) and L-glutamine (Sigma-Aldrich, UK). hBMSCs were used in passage 4. The mineralisation medium was comprised of basal culture media supplemented with 10 mM β-glycerophosphate (Sigma-Aldrich, Gillingham, UK), 50 μg/mL L-ascorbic acid (Sigma-Aldrich, Gillingham, UK) and 100 nM Dexamethasone (Sigma-Aldrich, Gillingham, UK). The culture medium utilised for EV isolation and dosing was depleted of FBS-derived EVs by ultracentrifugation at 120,000 g for 16 hr prior to use.

### GO-MNP synthesis and characterisation

GO-MNPs were synthesised as described elsewhere[17]. The synthesised GO-MNPs were functionalised with TREK1 antibody (Almone labs, APC-047) as mentioned previously[30]. The prepared TGMNPs were modified with 1μL of DOTAP (1,2-Dioleoyl-3-trimethylammonium propane) to enhance the internalisation and avoid particle agglomeration[31]. TGMNPs (25 µg) were added to cells and to provide magnetic stimulation to the cells, a custom-designed MICA bioreactor (MICA Biosystems, West Midlands, UK) was kept in an incubator maintained at 37[°C with 5% CO[. Additionally, control groups including those without MICA, without TGMNPs, and with TGMNPs alone were maintained under identical incubation conditions to study the MICA-mediated osteogenic inductive studies.

The X-ray diffraction (XRD) characterisation of MNPs and GO-MNPs was assessed using Bruker D2 phaser equipped with a cobalt source of wavelengths Ka1 1.7890 and Ka2 1.7929, step size 0.02028°. The Raman spectroscopy measurements were recorded using Renishaw inVia and the data was collected using a 633nm laser beam. The magnetic properties of the MNPs and GOMNPs were evaluated using a model 10vector VSM with a maximum field of 20kOe and sensitivity of 1×10^-6^emu at room temperature.

### MICA-induced osteogenesis

MC3T3 (4 x 10^4^) cells were seeded in osteogenic media in 24 well plates to assess the osteogenic potential of the TGMNPs. After 24 h, TGMNPs (25 µg) were added to each well and the cells were subjected to magnetic stimulation for 1h every day with a media change every 2 days. Following 3 and 7 daily treatments of magnetic stimulation, the ALP enzymatic activity was evaluated using the Sensolyte ALP assay kit. Briefly, the cells were lysed using 0.1 % Triton X. After 10 min, the cells were gently scraped and centrifuged at 2500g for 10 min at 4°C to separate cellular debris from the enzymatic supernatant. Afterwards, 50 µL of supernatant and 50 μl p-nitrophenyl phosphate solution was combined in each well of a 96-well plate and gently shaken for 30 s. Following 1h of incubation, the absorbance was measured at 405 nm using the microplate reader. The ALP concentration was then evaluated using a standard curve against known protein concentrations. Additionally, the total protein concentration was assessed using the BCA Protein Assay Kit (Thermo Scientific, USA).

### EV isolation

Osteoblasts were cultured at scale in 6 well plates (Sarstedt, UK) and the medium was collected every two days. TGMNPs (25 µg) were added to cells and were subjected to magnetic stimulation for 1h every day. EVs were isolated from the conditioned medium from the following groups: Untreated cells (CTL EVs), MICA stimulated cells (MICA EVs),

TGMNPs stimulated cells (TREK EVs) and MICA stimulated TGMNPs cells (MICA/TREK EVs). EVs were obtained from the conditioned medium by differential ultracentrifugation as previously described [32]: 2000 g for 20 min, 10,000 g for 30 min and 120,000 g for 70 min to pellet EVs. The supernatant was removed, and the pellet was washed in sterile PBS and centrifuged at 120,000 g for 70 min and the resultant pellet was re-suspended in 200 μl PBS. Ultracentrifugation was performed with the Sorvall WX Ultra Series Ultracentrifuge (Thermo Scientific, UK) and a Fiberlite, F50L-8×39 fixed angle rotor (Piramoon Technologies Inc., USA).

### EV Particle Size, Concentration and Tetraspanin Analysis

To determine the EV particle size, concentration and tetraspanin content, flow cytometry was conducted as previously described [33]. A NanoAnalyzer U30 (SPCM APDs) was used for the detection of side scatter (SSC) and fluorescence of individual particles. Measurements were taken over a 1-minute interval at a sampling pressure of 1.0 kPa, maintained by an air-based pressure module. Particle count was diluted to remain within the optimal range of 2000 -12,000/min.

The concentration of samples was determined by comparison to 250 nm silica nanoparticles of known concentration to calibrate the sample flow rate. EV isolates were sized according to standard operating procedures using a proprietary 4-modal silica nanosphere cocktail (NanoFCM Inc., S16M-Exo). Using the NanoFCM software (NanoFCM Profession V2.0), a standard curve was generated based on the side scattering intensity of the four different silica particle populations of 68, 91, 113 and 155 nm in diameter. The laser was set to 15 mW and 10% SSC decay.

To assess the EV tetraspanin phenotype, the following antibodies were used: FITC-conjugated anti-human CD63 (BioLegend), FITC-conjugated anti-human CD9 (Abcam, Cambridge, UK) and FITC-conjugated anti-human CD81 (Abcam, Cambridge, UK). EVs were diluted to 1 × 10^10^ particles/mL in PBS and 9 μL was mixed with 1 μL of conjugated antibody (single or mixed cocktail), before incubation for 30 min at room temperature. The incubation concentration ratio for single antibodies was 1:50 (in PBS) and 1:150 for the cocktail of 3 antibodies (1 μL of 1:5 of mixed antibody cocktail). After incubation, the mixture was diluted in PBS to 1 × 10^8^ -1 × 10^9^ particles/mL for analysis. Data processing was performed by the nFCM Professional Suite v2.0 software. The total EV protein concentration was determined using the Pierce Micro BCA Protein Assay Kit (Thermo Scientific, Paisley, UK).

### Transmission Electron Microscopy (TEM)

The MNPs, GOMNPs and EVs were imaged using the JEOL JEM1400 transmission electron microscope coupled with an AMT XR80 digital acquisition system. For EV, the vesicles were physisorbed to 200-mesh carbon-coated copper formvar grids (Agar Scientific, Stansted, UK) and 1% uranyl acetate was used for negative staining.

### EV-Induced hBMSC Osteogenesis

hBMSCs were seeded at a density of 21 × 10^3^ cells/cm^2^ in the basal medium within 96-well plates (Nunc, Thermo Scientific, Paisley, UK). After 24 h, the medium was replaced with a mineralisation medium supplemented with EVs derived from untreated (CTL EVs), MICA stimulated (MICA EVs), TGMNP only (TREK EVs) and MICA with TGMNPs stimulated osteoblasts (MICA/TREK EVs) (10 μg/mL) for 14 days. The EV-supplemented mineralisation medium changes were performed every 48 hrs. Cells cultured in the mineralising medium alone were used as the control.

### Collagen Production

Picrosirius red staining was conducted to assess extracellular matrix collagen production. Briefly, cells were washed twice in PBS, fixed in 10% NBF for 30 min, and then stained with 0.1% Sirius red in saturated picric acid (Sigma-Aldrich, Gillingham, UK) for 60 min. Acetic acid (0.5 M) wash was used to remove the unbound dye, followed by a distilled water wash. Samples were left to air dry prior to imaging using light microscopy (EVOS XL Core, Invitrogen, Paisley, UK).

### Calcium Deposition

Alizarin red staining was used to evaluate the extracellular matrix calcium deposition. Briefly, cells were washed twice in PBS and fixed in 10% NBF for 30 min. Samples were washed in distilled water and incubated with alizarin red solution (Sigma-Aldrich, Gillingham, UK) for 10 min. Distilled water was used to remove the unbound dye. Calcium deposition was visualised using light microscopy (EVOS XL Core, Invitrogen, Paisley, UK).

### LC-MS Sample Preparation and Analysis

Samples were heated to 95 °C for 5 minutes followed by sonication. Protein was extracted through the addition of 400 µL of ice-cold acetone and incubated at -80 °C for 1 hour before centrifugation at 14,000 xg for 10 minutes. Supernatant was discarded and the pellets air dried. A 0.1% RapiGest (Waters Corporation, Milford, MA, USA) solution was prepared in 50 mM Ammonium Bicarbonate (pH 7.8). 50 µL was added to each pellet prior to incubating at 80 °C for 45 minutes, followed by centrifugation at 14,000 xg for 10 minutes. Dithiothreitol (DTT, 5 mM) (Fisher Scientific, Loughborough, UK) was added and to the samples, which were heated to 65°C for 20 minutes for protein denaturation. Once cooled, 15 mM Iodoacetamide was added at room temperature and placed in the dark for 30 minutes. This was followed by incubation in 1 µg trypsin (ThermoFisher Scientific, UK) overnight at 37°C. Samples were acidified (0.5% v/v formic acid) and incubated at 37°C for 25 minutes. Finally, samples were centrifuged at 21,000 xg for 20 minutes and the supernatant collected and stored at −80°C for liquid chromatography mass spectrometry (LC-MS) analysis.

The ACQUITY M Class (Waters Corporation, Milford, MA, USA) with a Symmetry C18 5 μm, 2 cm × 180 μm pre-column and a High Strength Silica (HSS) T3 C18 1.7 μm, 15 cm × 75 μm analytical reversed-phase column (Waters Corporation, Milford, MA, USA) was utilised to perform one-dimensional nanoscale LC separation of tryptic peptides. The analytical column temperature was set to 35^◦^C. MBV samples were transferred to the pre-column at 15 μL/min for 2 minutes with mobile phase A; aqueous 0.1% (v/v) formic acid. Peptides were eluted and separated with a gradient of 3%–40% of mobile phase B (acetonitrile with 0.1% (v/v) formic acid) for 90 minutes at 400 nL/min. Lock mass solution was delivered to the reference sprayer at 1 μL/min by the LC system auxiliary pump and sprayed with a frequency of 60 s. Mass spectrometric analysis was acquired using the SELECT SERIES™ Cyclic Ion Mobility Mass Spectrometer (Waters Corporation, Wilmslow, UK) in v-mode with a nominal resolution of 35,000 full width at half maximum (FWHM) in positive mode electrospray ionization (ESI). The ion source block temperature was at 100°C and capillary voltage at 3.2 kV. The time-of-flight analyser was externally calibrated with NaCsI from m/z 50 to 1990. The data were post-acquisition lock mass-corrected using the doubly charged monoisotopic ion of (Glu1)-Fibrinopeptide B (m/z 785.8426). Accurate mass LC-MS data were collected in a randomised order using the ion mobility-enabled, data-independent acquisition mode (HDMSE) for 0.5 seconds with a 0.02 second interscan delay. A low and elevated energy data cycle was acquired each second, where transfer collision energy was 6 eV (per unit charge) in low energy mode and was increased from 19 to 45 eV (per unit charge) in 0.5 seconds in elevated energy mode.

### Data Processing and Bioinformatics

Progenesis QI for Proteomics version 4.2 (Nonlinear Dynamics, Newcastle upon Tyne, UK) was used to process all acquired data. Protein identifications were obtained by the reviewed entries of a murine UniProt database (20,405 reviewed entries, release 2022_12). To detect and monitor protein and peptide identification error rates (1% FDR), decoy database strategies were utilised. Peptide and fragment ion tolerances were determined automatically, one missed cleavage site was allowed, as well as fixed modification carbamidomethylation of cysteine. Variable modifications were also specified, which included the oxidation of methionine and deamidation of asparagine and/or glutamine. From the abundance data obtained by Progenesis, linear regressions were plotted using Origin Lab 2020. The protein annotation through evolutionary relationship (PANTHER) classification systems (version 19.0) was used for gene ontology (GO) annotation of biological pathways, molecular mechanisms and cellular components of protein found to be significantly upregulated in MICA/TREK EVs. StringDB was used to generate a protein-protein interaction network of differentially expressed proteins [34].

### Statistical analysis

For all data, experiments were performed in triplicate. Statistical analysis was assessed using the IBM SPSS software (IBM Analytics, version 21). The Shapiro[Wilk test was used to analyse the normality of data. Data that was proven to be normally distributed were analysed using parametric students’ T[test, one[way ANOVA, or paired T[test. Non[normally distributed data were assessed using non[parametric Mann[Whitney t[test or Kruskal[Wallis ANOVA. P values equal to or lower than 0.05 was considered as significant.

*P ≤ 0.05, **P ≤ 0.01 ***P ≤ 0.001.

## 3. Results and Discussion

### 3.1. Characterisation of Graphene Oxide Magnetic particles

Previous studies have shown how GO demonstrates enhanced biocompatibility and surface area in comparison to pure MNPs, with higher functionalisation potential and excellent electrochemical properties[17] making it an ideal coating material for use with magnetic particles. GO-MNPs offer superior TREK1 binding efficiency due to their increased surface area, π-π stacking interactions and the presence of functional groups such as hydroxyl and carboxyl groups that can improve protein anchoring[17, 35]. These characteristics make GO-MNPs a promising platform for targeted mechanobiology applications.

Monodispersed MNPs with uniform size (∼25nm), shape and crystallinity were synthesised using the high-temperature thermal decomposition method as previously reported[17] and seen in the TEM image shown in Figure 2A. MNPs were amine functionalised with APTES through a ligand exchange mechanism and coupled with the carboxyl groups of GO sheets through an EDC/NHS binding followed by functionalisation with TREK1 antibody to enable the selective targeting. The morphology of the discreet GO-MNPs, shown in the TEM image (Figure 2B) clearly indicates the uniform distribution of MNPs over the GO surface and a distinct view of GO sheets along the border of the GO-MNP composite.

**Figure 2.**
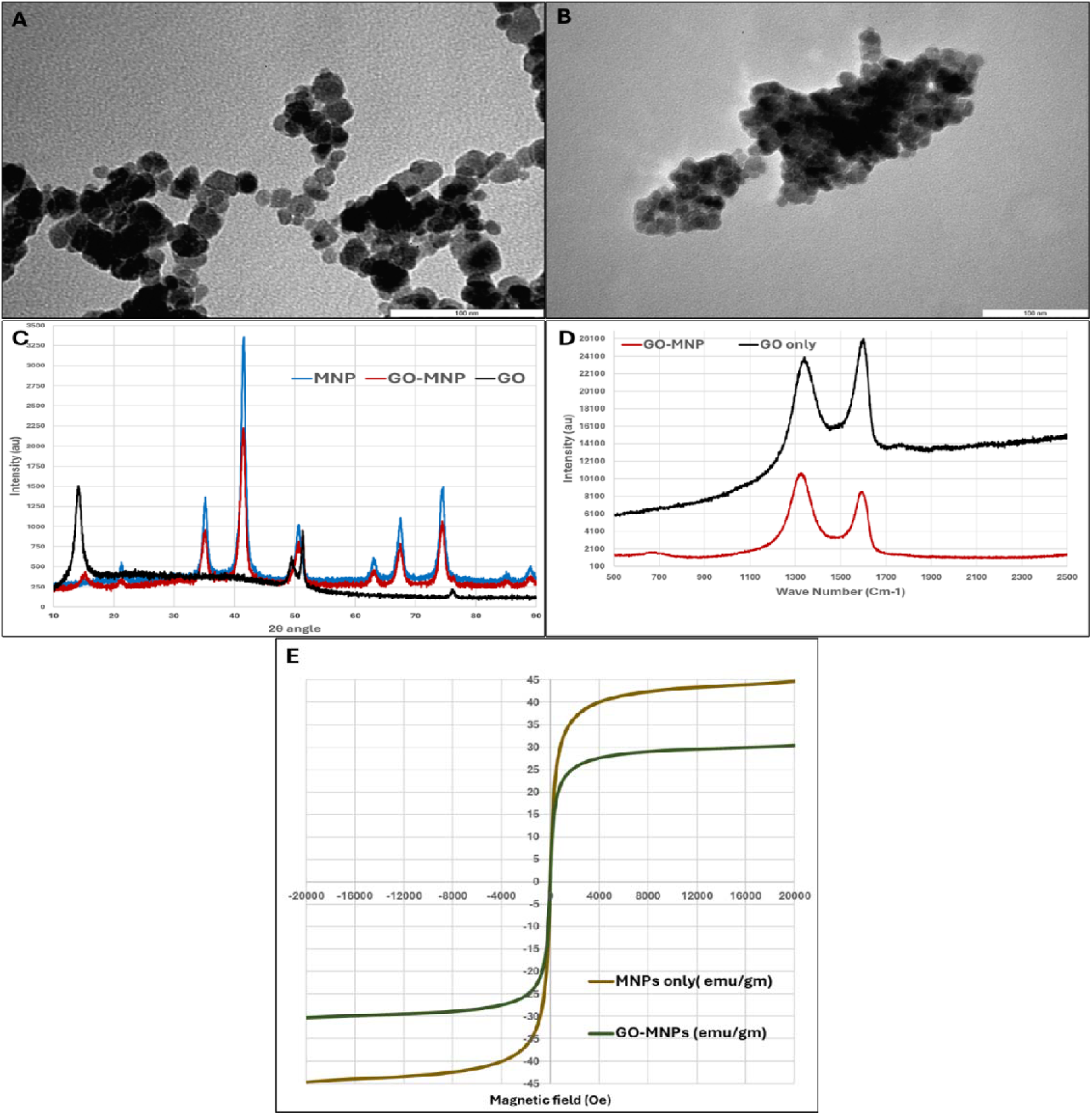
Characterisation of MNPs and GO-MNPs, TEM images of A) MNPs, B) GO-MNPs, C) XRD analysis, D) Raman spectroscopy, and E) VSM measurement of magnetic hysteresis loops of MNPs and GO-MNPs.

The crystallinity and phase composition of the MNPs, GO and GO-MNPs were investigated through the XRD analysis. The XRD pattern of the synthesised MNPs showed the characteristic XRD diffraction peaks at 2θ= 21.3°, 35.2°, 42.5°, 51.2°, 56.9°, 62.0° and 74.5° indicating the cubic inverse spinel structure of Fe_3_O_4_ (Figure 2C). Meanwhile, GO showed the characteristic XRD peak around 2θ= 12 ° indicating the (002) plane with an interlayer distance of approximately 0.84nm. This increased interlayer distance is an indication of oxygen-containing functional groups such as hydroxyl and carboxyl groups on the GO[36, 37]. The obtained XRD pattern is in accordance with the previously reported values hence confirming the successful synthesis of GO [38]. The XRD spectra of GO-MNP (Figure 2C) showed all the key peaks of both MNP and GO confirming the formation of a highly crystalline GO-MNP composite with MNPs well integrated into the GO sheets. Moreover, the absence of any unwanted peaks indicates the purity of the synthesised GO-MNPs with no detectable secondary phases or byproducts. The peak broadening observed in the GO-MNP compared to MNPs and GO also indicates a nanoscale crystallite size further confirming the uniform distribution of MNPs on GO sheets.

Raman spectroscopy is a powerful tool for characterising the structural and electronic properties of GO. The Raman spectra of GO revealed two prominent peaks at 1340 cm^-1^ and 1601 cm^-1^ indicating D-band and G-band respectively. Here the D-band at 1340 cm^-1^ is associated with the defects and disorder in the sp^2^ carbon network and the G-band at 1601 is linked to the in-plane stretching of sp^2^ carbon bonds[39]. As seen in Figure 2D, the Raman spectra of GO-MNP showed peaks around 1331 cm^-1^ (D-band) and 1597 cm^-1^ (G-band) indicating the presence of both MNPs and GO in the GO-MNP composite. The observed peak shifts in GO-MNP from the GO-only sample in the D-band indicated the possible charge transfer between GO and MNPs[40]. Similarly, the shift in the G-band suggests the influence of MNP-induced strain or electronic interactions affecting the GO-structure further confirming the formation of GO-MNP composite while retaining the basic structural integrity of GO[41].

The superparamagnetic behaviour of GO-MNP can be validated through the VSM as it is essential for the controlled and non-invasive magnetic stimulation of mechanosensitive ion channels. The hysteresis loop revealed that both MNPs and GO-MNPs possess a superparamagnetic behaviour with no evidence of coercivity and remanence (Figure 2E). The saturation magnetisation values of MNPs and GO-MNPs were observed to be 44.7 emu/g and 30.4 emu/g respectively. The observed lower saturation magnetisation for GO-MNP can be due to the presence of GO sheets which are non-magnetic in nature. This non-magnetic nature of GO may interfere with the magnetic alignment of MNPs, thereby leading to a decrease in the magnetic saturation of the GO-MNP composite.

### 3.2. MICA/ TGMNPs-induced osteogenic differentiation of osteoblasts

To evaluate the osteoinductive potential of the TGMNPs under MICA stimulation, ALP activity was measured as a key early-stage marker of osteogenesis. After 3 and 7 days of treatment with TGMNPs, ALP activity in the MICA-treated groups was significantly enhanced compared to non-MICA groups at days 3 and 7 (Figure 3A). Additionally, the ALP on day 7 is higher than that of day 3. The obtained results signify the enhanced osteogenic potential of the MICA-treated groups, highlighting their ability to promote early-stage osteoblast differentiation more effectively than the non-MICA treatments.

**Figure 3.**
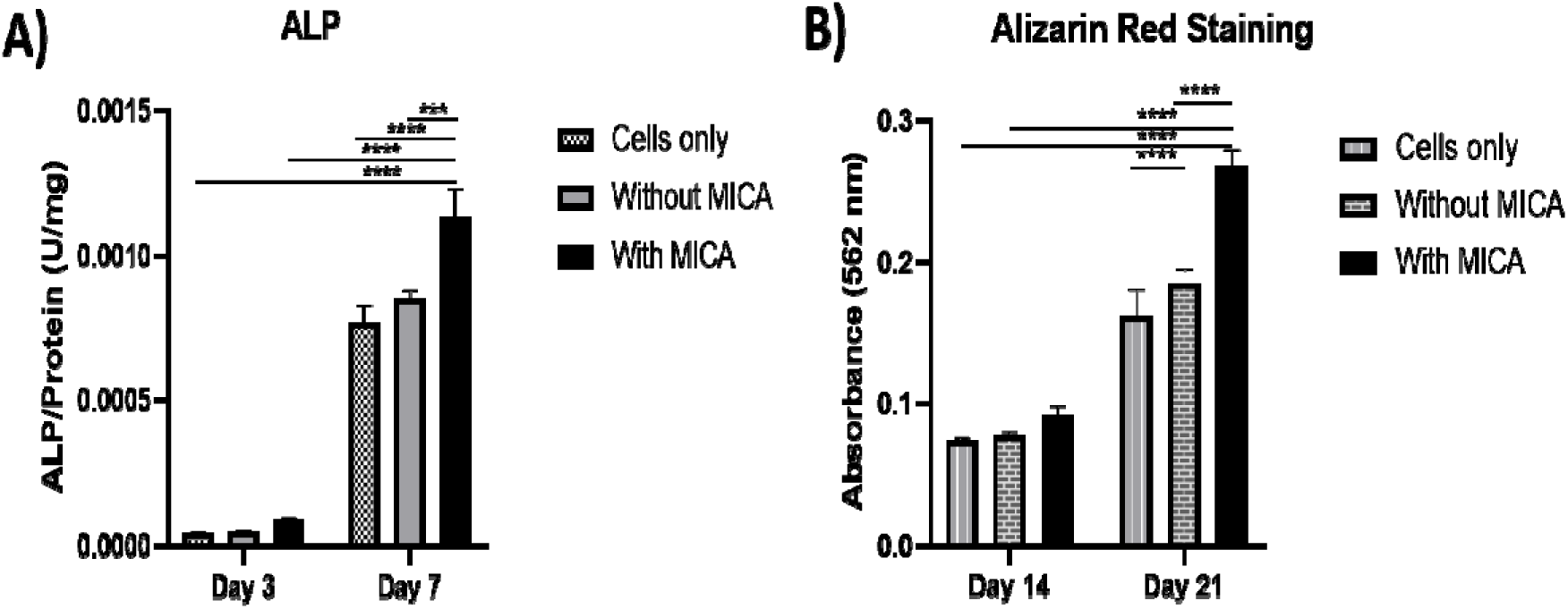
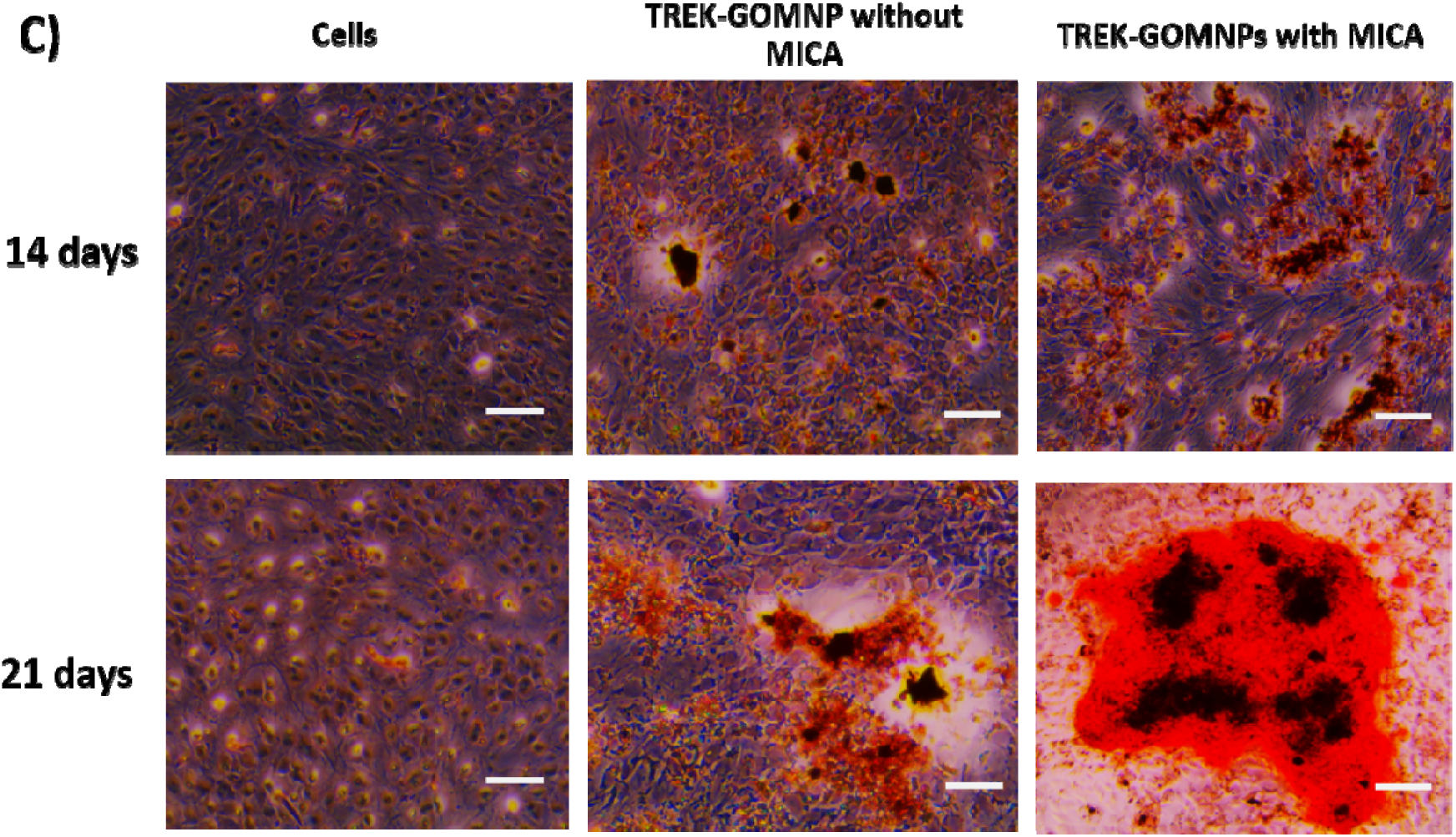
Osteogenic differentiation potential of MICA /TGMNPs using MC3T3 cells. A) ALP activity after 3 and 7 days culture with/without MICA stimulation. B) Quantitative analysis of Alizarin red staining using CPC after 14 and 21 days of treatment with/without MICA. C) Representative optical microscopy images of MC3T3 after Alizarin Red staining (Scale 200 µm).

Alizarin Red staining was performed to assess extracellular matrix mineralisation, a hallmark of late-stage osteogenesis, in both MICA and non-MICA-treated groups after 14 and 21 days of treatment. Our findings showed that TGMNPs treatment alone increased MC3T3s calcium deposition when compared to the untreated control (Fig 3B, C). MICA stimulation in conjunction with TGMNPs treatment further enhanced the mineralisation potential of these cells compared to TGMNPs alone and the untreated control, consistent with the ALP activity results.

#### Isolation and characterisation of MICA/TREK-EVs

EVs were then isolated from the conditioned medium of osteoblasts tagged with TGMNPs and stimulated with/without MICA over a 2-week culture period using differential ultracentrifugation. EVs were isolated from the conditioned medium from the following groups: Untreated cells (CTL EVs), MICA stimulated cells (MICA EVs), TGMNPs stimulated cells (TREK EVs) and MICA stimulated TGMNPs cells (MICA/TREK EVs). TEM imaging showed the obtained EVs exhibited a typical spherical morphology indicative of nano-sized vesicles (Figure 4A), consistent with previous studies [32]. The nano-flow cytometry analysis detected particles with an average diameter of 65.32 ± 0.46 nm, 71.36 ± 0.60 nm, 71.53 ± 0.15 nm, and 67.92 ± 0.22 nm for the CTL EVs, MICA EVs, TREK EVs and MICA/TREK EVs respectively (Figure 4B), corroborating with the TEM images.

**Figure 4.**
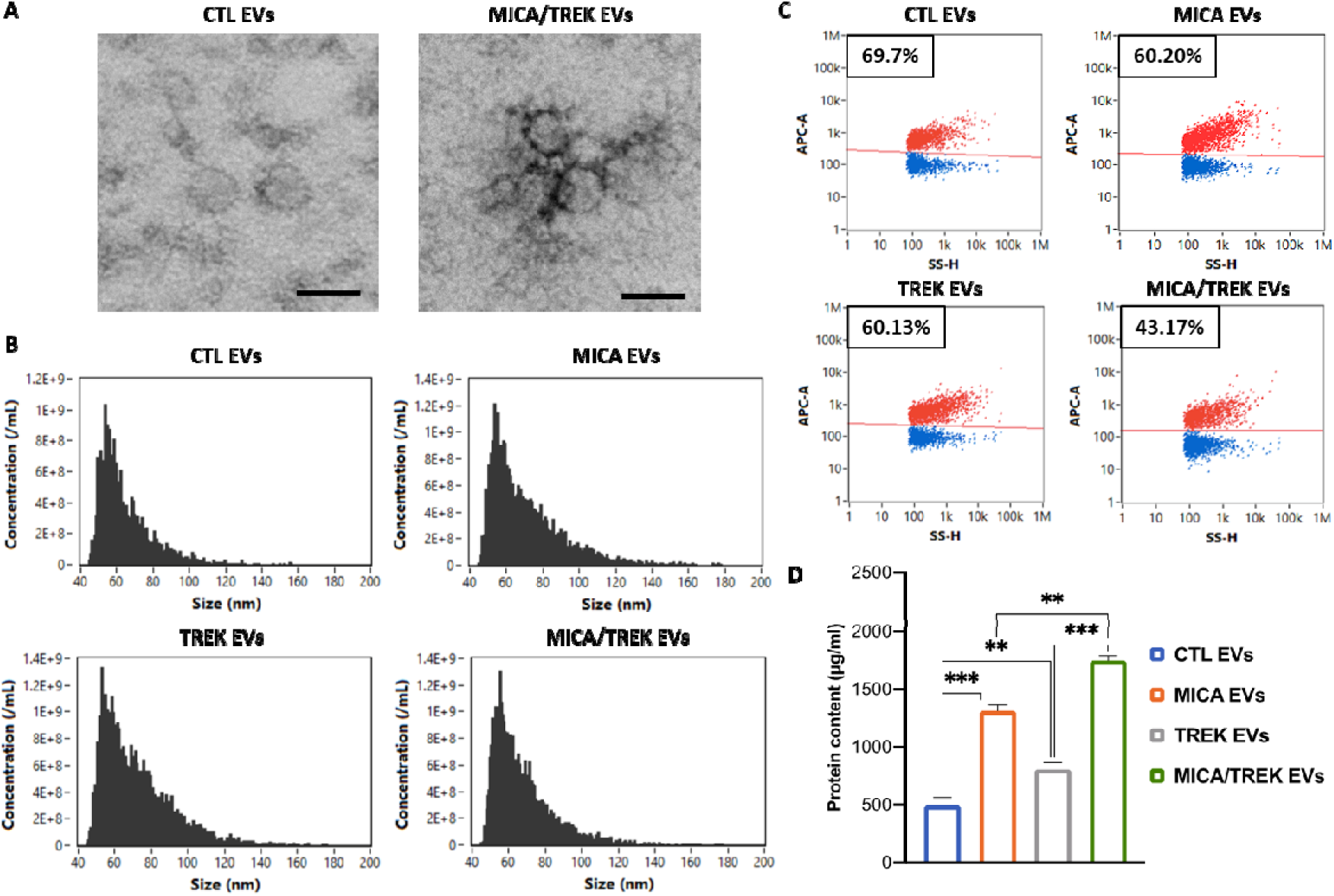
Characterization of EVs isolated from MICA/ TGMNPs stimulated and untreated mineralising osteoblasts. A) TEM images of EVs obtained from MICA/TREK stimulated and untreated osteoblasts. Scale bar = 50 nm, B) Flow cytometry analysis depicting the size distribution of particles. C) Single-particle phenotyping of osteoblast-derived EVs. EVs were fluorescently labelled with APC-conjugated antibodies specific to CD9, CD63 and CD81. D) Protein quantification of isolated EVs. Data are expressed as mean ± SD (n = 3).

The positive staining percentages of CD9, CD63 and CD81 for the CTL EVs were 66.63%, 36.77% and 27.77%; MICA EVs were 55.70%, 40.97% and 31.33%; TREK EVs were 57.00%, 39.80% and 32.57%; MICA/TREK EVs were 32.63%, 27.13% and 15.50%. When assessing triple-positive staining, 69.17%, 60.20%, 60.13% and 43.17% of all particles stained positive for the CTL EVs, MICA EVs, TREK EVs and MICA/TREK EVs respectively (Figure 4C).

EV protein quantification showed that EVs acquired from the MICA/TREK, MICA, and TREK stimulated groups exhibited a 3.53-fold, 2.65-fold, and 1.62-fold enhanced protein content when compared to EVs obtained from the untreated cells (Figure 4D) (P < 0.001). These findings confirmed that magnetic stimulation and the addition of TGMNPs during osteoblast mineralisation enhanced the EV protein yield during culture. The influence of mechanotransduction on EV production yield has been reported in the literature. For example, the use of 3D-printed bone-mimetic titanium scaffolds significantly enhanced the production of osteoblast-derived EVs when compared to cells cultured on tissue culture plastic [42]. Mechanotransduction induced by hydrostatic pressure significantly increased the production of EVs from cartilage microtissues [43]. Importantly, our data showed superior EV production yield combining the synergistic effects of magnetic stimulation and TGMNPs, highlighting the potential of harnessing this platform technology for the scalable manufacture of EVs.

### 3.3. MICA/TREK-EVs Enhanced hBMSCs Extracellular Matrix Mineralization

In this study, hBMSCs were treated with osteoblast-derived EVs manufactured under MICA/TREK stimulation. Extracellular matrix production was assessed by Picrosirius Red staining to evaluate collagen production. Picrosirius red staining can selectively bind to collagen fibers thus enabling the visualisation of extracellular matrix formation [44, 45]. Our findings showed EVs acquired from MICA/TGMNPs stimulated osteoblasts exhibited greater hBMSCs collagen production when compared to those cells treated with EVs from MICA, TGMNPs, or untreated groups (Figure 5A). The extent of extracellular matrix mineralisation was evaluated via Alizarin Red staining to detect calcium-rich deposits. Our findings show that MICA/TREK-EV treatment substantially increased calcium deposition when compared to the treatment with EVs derived from the MICA, TREK, or untreated groups (Figure 5B), consistent with the collagen production results. Previous studies demonstrated that MICA/TREK stimulated MG63 osteosarcoma cells and immortalised MSCs mineralisation [46], suggesting the influence of MICA/TREK on enhancing the production of osteoinductive EVs propagating the enhanced osteogenic phenotype.

**Figure 5.**
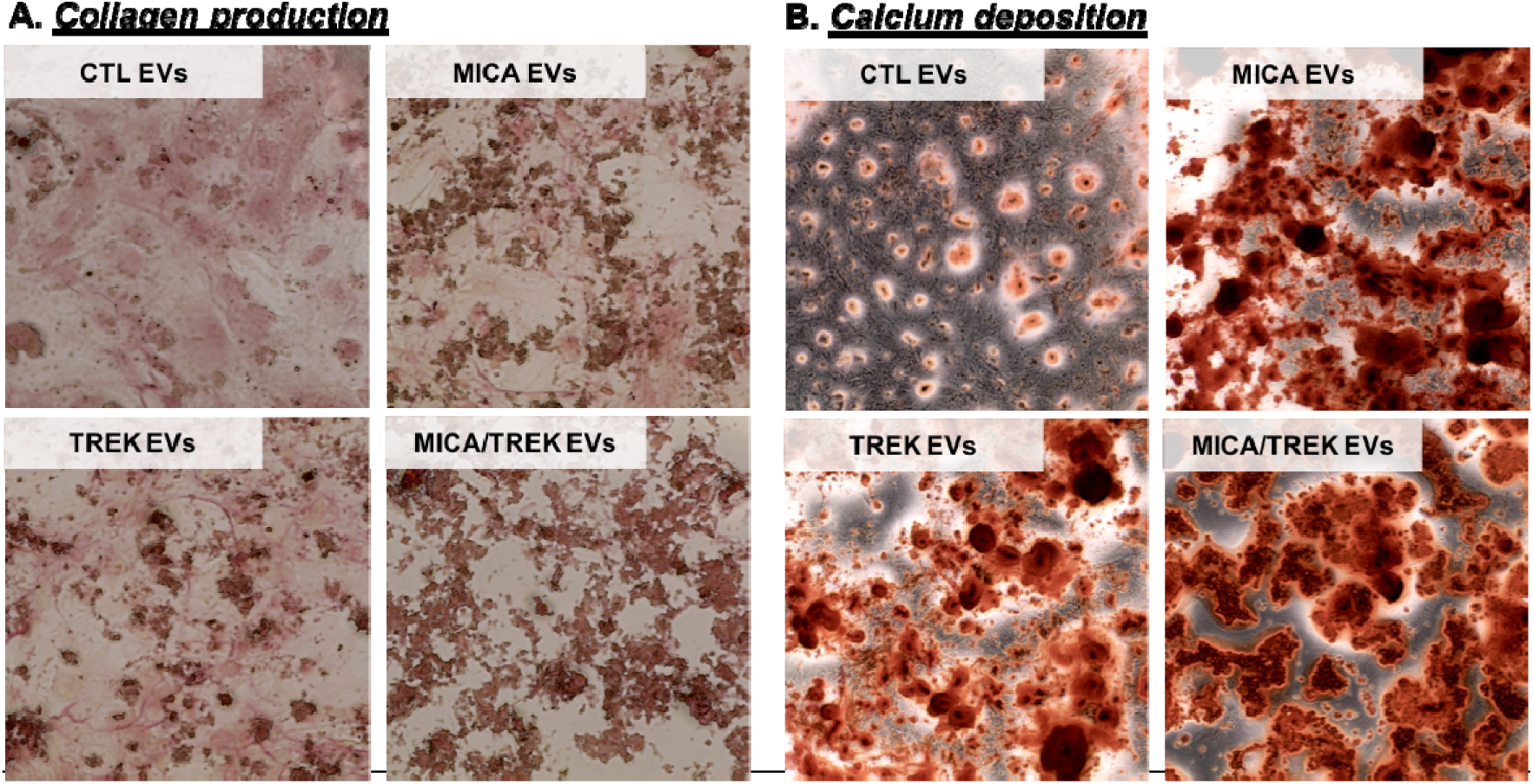
Osteogenic differentiation of hBMSCs treated with osteoblast-derived EVs manufactured with/without MICA/TREK stimulation. A) Picrosirius red staining for collagen production, and B) Alizarin red staining for calcium deposition following 14 days of osteogenic culture.

**Figure 6.**
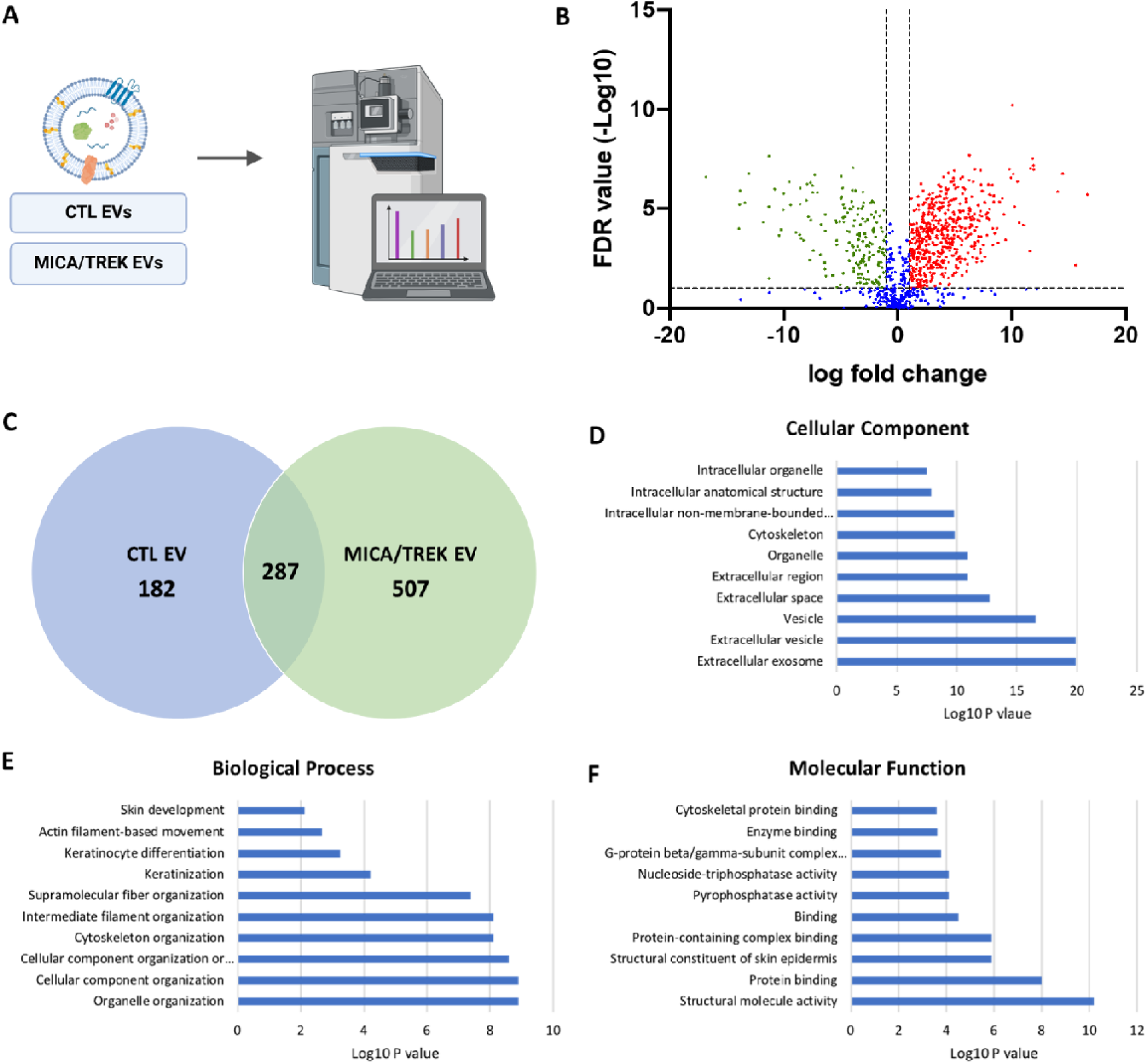
A) Analysis of differentially expressed proteins from CTL EVs and MICA/TREK EVs. B) Volcano plot displaying Log2 values for the proteins fold-change against Log10 FDR. Proteins with a Log2 fold difference below 1 and a statistical value of > 0.05 were not considered to be statistically significant (vertical and horizontal lines respectively). The red points in the plot represent the significantly upregulated MICA/TREK EV proteins, and the green points represent significantly upregulated CTL EV proteins. C) Venn diagram comparing proteins differentially expressed from EVs. A total of 287 shared proteins; 507 proteins upregulated in MICA/TREK EVs and 182 proteins upregulated in CTL EVs. Top ten GO prediction scores covering the domains of D) cellular components, E) biological processes and F) molecular function of proteins significantly upregulated in MICA/TREK EVs.

**Figure 7.**
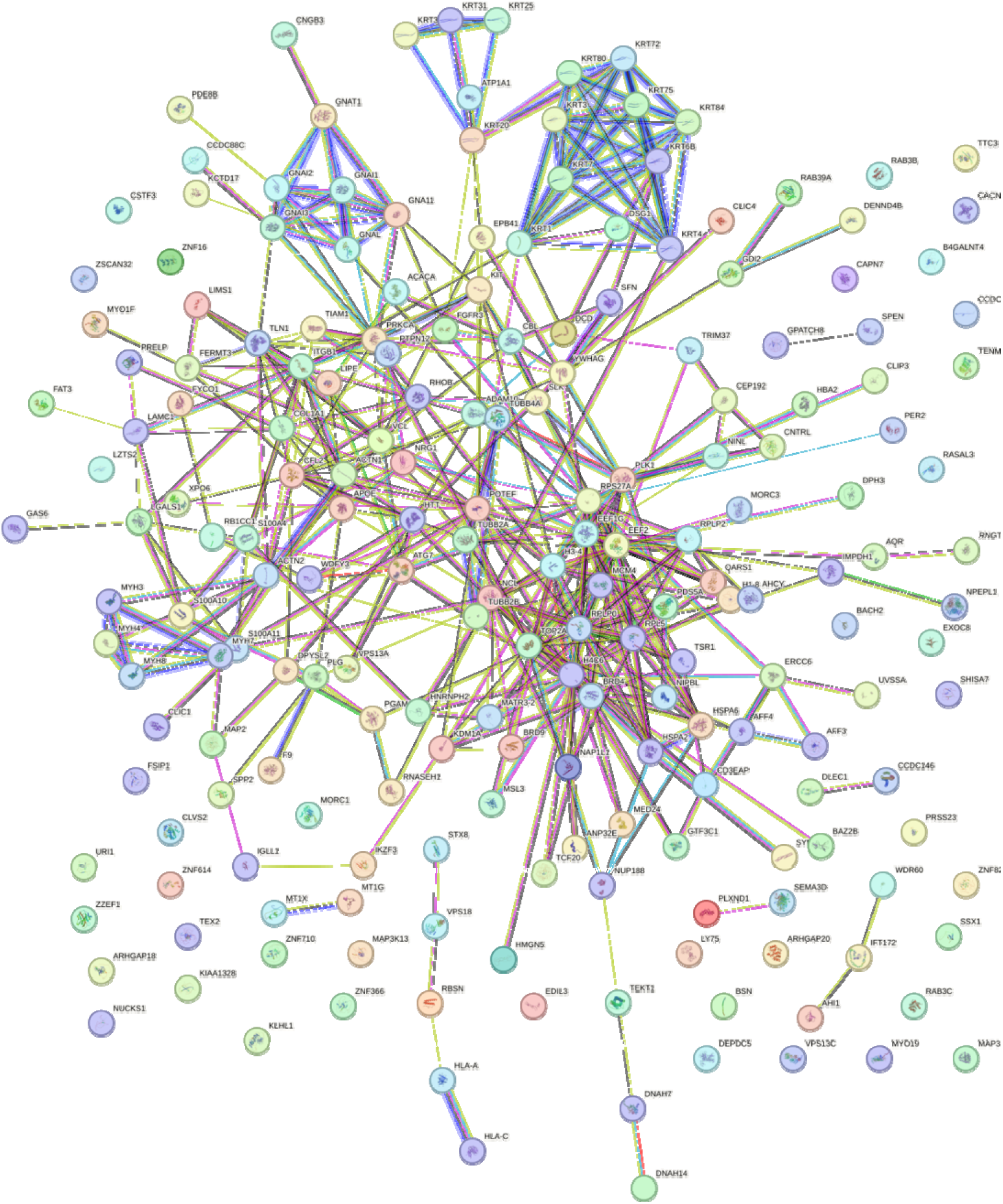
String DB network illustrating interactions between proteins in the MICA/TREK EVs, with a significant degree of protein-protein interaction (p < 10-16).

### 3.4. Proteomics analysis of MICA/TREK EVs

Having confirmed the striking improvement of synergistic MICA/TREK stimulation of osteoblast EV osteoinductive potential, vesicle protein content was profiled to further elucidate their possible mechanism of action. The proteomes of the CTL EVs and MICA/TREK EVs were compared for three independent sample preparations using a label-free MS-LC/LC approach. The use of stringent criteria only permitted the inclusion of proteins identified in a least two biological replicates, with > 2 spectral counts in at least one repeat. The volcano plot shows the variation of protein expression between the untreated and MICA/TREK EVs. Protein database searching resulted in the identification of a total of 995 proteins. Of these, 507 proteins were significantly upregulated in MICA/TREK EVs, 182 upregulated in CTL EVs, and 287 shared proteins. The differentially expressed proteins are identified in Table S1. To provide an overview of the principal processes, mechanisms and cellular locations of proteins significantly upregulated in MICA/TREK EVs, GO analysis was performed. Significantly enriched MICA/TREK EVs proteins were found to be associated with GO functional annotation of molecular functions (i.e. structural molecule activity, protein binding), cellular components (i.e. extracellular vesicles, cytoskeleton) and biological processes i.e. organelle organization, cytoskeleton organization). A String DB network was constructed to further investigate any potential interactions between proteins associated with EVs (Figure X) revealing a significant degree of protein-protein interaction (P[<[10-16).

Our findings showed that the MICA/TREK EVs were enriched with pro-osteogenic proteins. Among them were the calcium channelling annexin proteins, where several proteins of this family were significantly upregulated within the MICA/TREK EVs (i.e. Annexin A2, A4, A5, A6, A11). These transmembrane proteins are known to play critical roles in the binding to the extracellular matrix [47, 48]. Moreover, these proteins are involved in transporting extracellular calcium into the lumen of EVs, resulting the mineral nucleation and ECM mineralization [49]. This highlights the possible role of the MICA/TREK EVs in stimulating recipient hBMSCs ECM mineralization observed in this study. Moreover, it provides indications of the role of MICA/TREK stimulation in producing EVs with superior ECM binding and mineralization potential.

The proteomics analysis identified the enrichment of several Rab proteins within the MICA/TREK EVs. Rab proteins are small GTPases that act as key regulators of intracellular membrane trafficking, from the formation of transport vesicles to their fusion with membranes [50, 51]. This function is essential for osteoblasts to produce and secrete the materials needed for bone formation [52]. RAB1A regulates vesicular protein transport from the endoplasmic reticulum (ER) to the Golgi compartment and onto the cell surface and plays a role in IL-8 and growth hormone secretion [53]. Moreover, it was reported that inhibiting Rab32 with the miR-124a, impaired EV secretion in lung cancer cells [54]. Studies have also suggested the role Rab proteins play in coordinating early osteogenesis [55]. Thus, the upregulation of several members of the Rab family within MICA/TREK EVs, indicates the influence of MICA on stimulating intracellular membrane trafficking, likely contributing to the enhanced EV production and osteoinductivity observed in this study.

Heat shock proteins (HSPs) play a crucial role in bone formation. Primarily known for their role in protecting cells from stress, HSPs also act as molecular chaperones, assisting in the proper folding and assembly of proteins essential for bone development [56–58]. Particularly, the MICA/TREK EVs were significantly upregulated with the HSP70 protein. Several studies have reported the influence of HSP70 in regulating osteogenesis. Li et al. highlighted the importance of HSP70 in MSC osteogenic induction, where following HSP70 knockdown, they observed significantly reduced MSC osteogenic marker expression [59]. Chen et al. conducted microarray and pathway analyses revealing that HSP70 promotes MSC osteogenesis via the activation of the ERK signalling pathway [60]. Moreover, studies have reported the enrichment of HSP within EVs during bone homeostasis [61]. Thus, the MICA-induced cellular stress likely upregulated the production of HSP70 in MC3T3s, leading to their enrichment within secreted EVs for autocrine/paracrine signalling.

It has been reported that calcium signalling plays an important role in the synthesis and release of EVs. Our findings show that the MICA-TREK EVs were enriched with the Voltage-Sensitive Calcium Channel (VSCC) protein. VSCCs have been shown to play an important role in bone cell regulation and are important regulators of intracellular calcium signalling in skeletal tissues[62]. The key role of VOCCs in mechanotransduction has been elucidated and it has been shown that mechanical force induces Ca^2+-^dependent contractions of the osteocyte cell membrane mediating EV release and demonstrates that EVs are another mechanism by which VSCCs influence the secretion of bioactive molecules within bone [62, 63]. EV signalling plays an important role in regulating bone remodelling during mechanical stimulation [64]. Considering the role that VSCCs have in differentiation and mechanically induced responses, Ca^+2^ influx via VSCCs could modulate EV secretion in the skeleton to regulate bone remodelling. Thus, the enrichment of VSCC proteins in MICA/TREK EVs indicates the role of these nanoparticles in stimulating Ca^2+^ signalling and subsequent EV release in recipient hBMSCs.

Proteomics analysis also highlighted the enriched transcriptional regulating proteins within the MICA/TREK EVs. There has been growing evidence regarding the role of epigenetic regulation in controlling lineage-specific differentiation [65, 66]. Our findings show that the MICA/TREK EVs were enriched with the epigenetic modifying protein histone acetyltransferase - KAT6A. This protein is involved in the addition of acetyl groups to histone proteins, ultimately enhancing the chromatin’s transcriptional activity [67]. Studies have shown that increasing acetylation within MSCs through the inhibition of histone deacetylase (HDAC) by HDAC inhibitors, significantly enhances osteogenic differentiation [68–70]. Previous studies have reported that HDAC inhibition in osteoblasts significantly enriched their EVs with epigenetic modifying proteins and microRNAs, which contributed to enhancing the vesicle’s osteoinductive potency [32]. Moreover, it has been reported that KAT6A plays a crucial role in maintaining the stemness of MSCs through regulation of the Nrf1/ARE signalling pathway, thus inhibiting ROS accumulation [71]. The MICA/TREK EVs were also enriched in lysine-specific histone demethylase 1A - KDM1A. Lysine demethylases control the process of histone methylation, which contributes to controlling chromatin structure and gene expression [72]. Studies have reported the importance of KDM1A in osteoblast differentiation, where Rummukainen et al. reported that KDM1A knockdown in MSCs led to a reduction in osteoblast activity and disrupted bone formation *in vivo* [73]. Moreover, the induction of hypomethylation within MSCs resulted in the production of EVs with enhanced osteoinductive potency [33]. Together, these findings demonstrate that MICA/TREK stimulation significantly enriched osteoblast EVs with epigenetic regulating proteins, which likely contributed to augmenting the transcriptional activity in recipient hBMSCs, enhancing osteogenic differentiation.

It is important to note that due to the diverse biological cargo of EVs, is it likely that the osteoinductive capacity of MICA/TREK EVs is a combination of changes across all EV components (i.e. metabolites, lipids, proteins, RNA species etc.), although this would require further investigation.

## 4. Conclusion

In conclusion, these findings demonstrate that enhancing the osteogenic differentiation of osteoblasts via MICA stimulation, significantly increased EV production yield and their osteoinductive potency. Furthermore, proteomics profiling revealed that the MICA/TREK EVs were enriched with proteins involved in ECM binding, osteogenic differentiation, EV signalling, and transcriptional regulation. These findings showcase the considerable utility of harnessing MICA as a novel engineering approach to enhance the scalable production of EVs as an acellular tool for bone repair. To our knowledge, this is the first study to promote the production yield and therapeutic potency of EVs for bone regenerative strategies through MICA.

## Acknowledgements

This research has been funded by ERC DYNACEUTICS (789119) Remote control healing: Next generation mechano-nano-therapeutics

## Data Availability Statement

The data that support the findings of this study are available from the corresponding author upon reasonable request.

## Supplementary information

**Supplementary Table 1.**
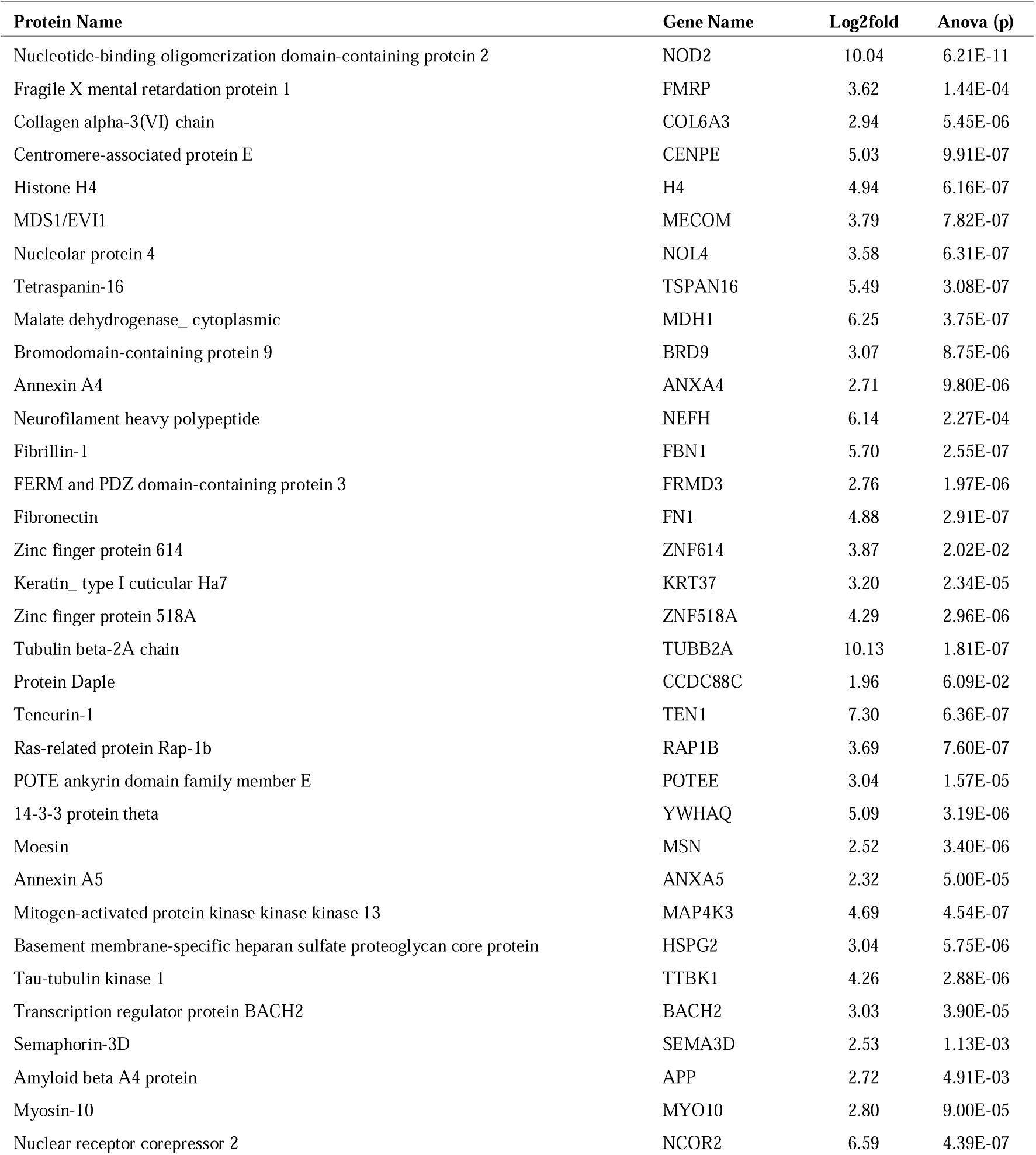
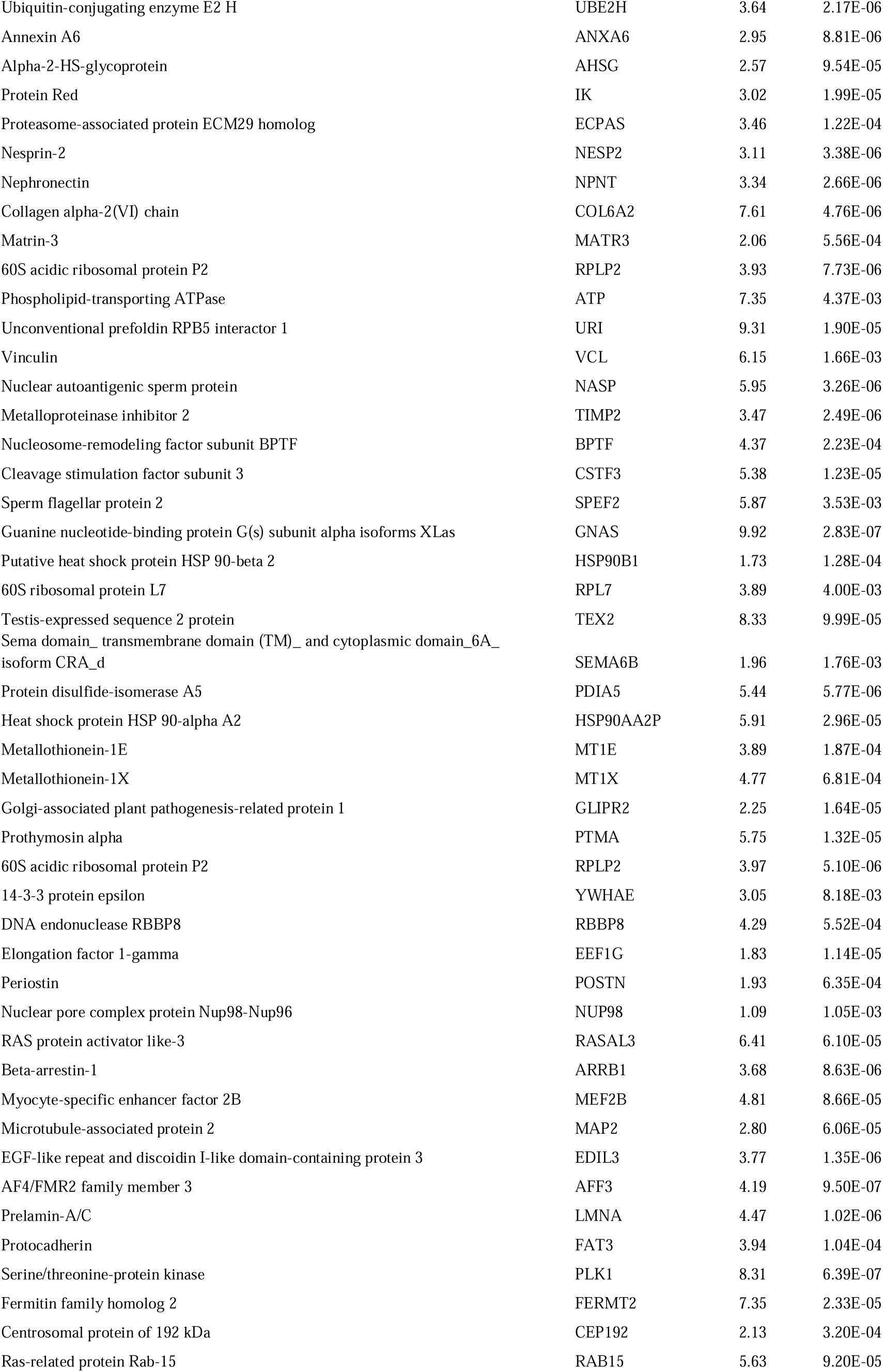
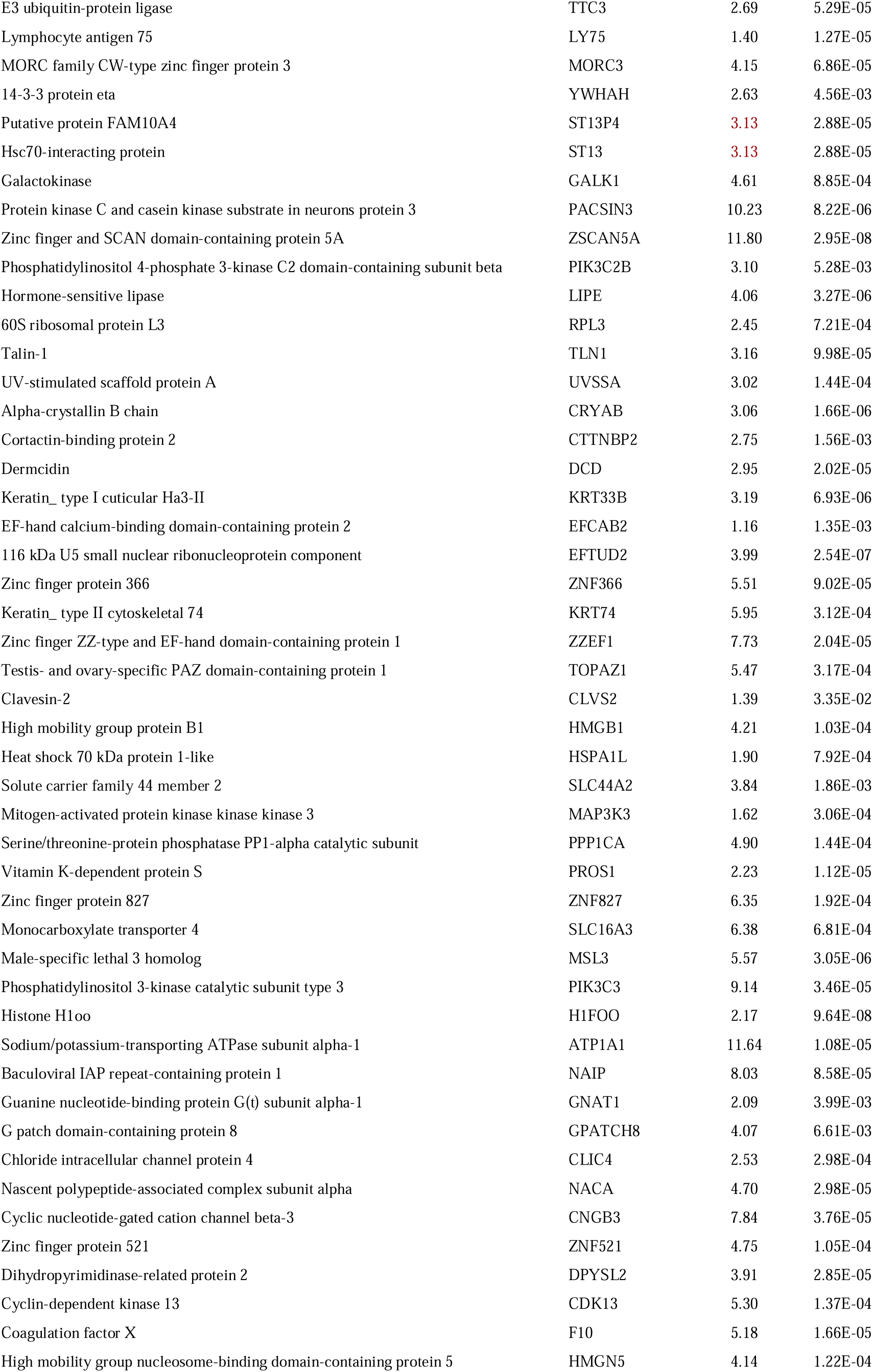
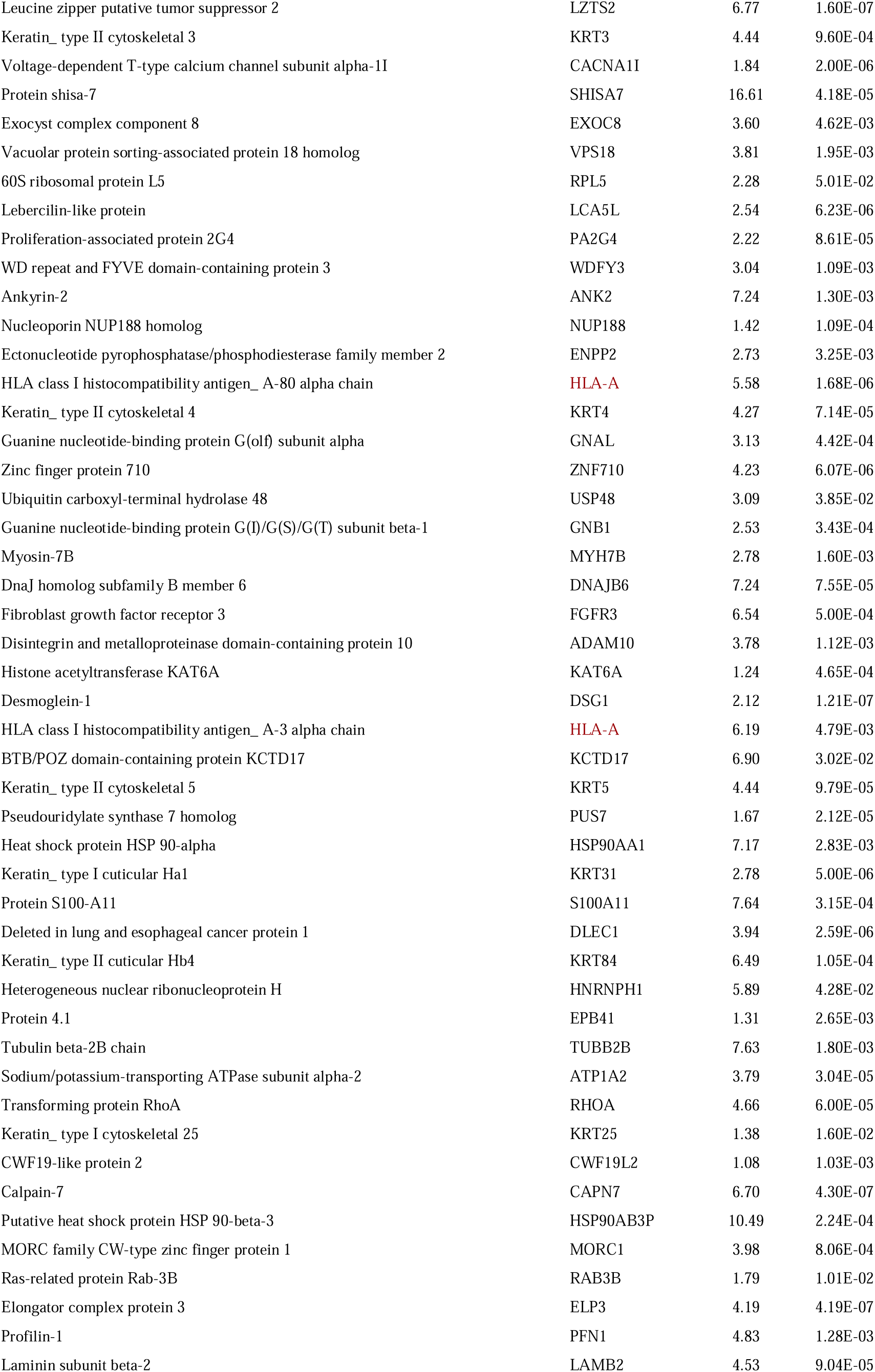
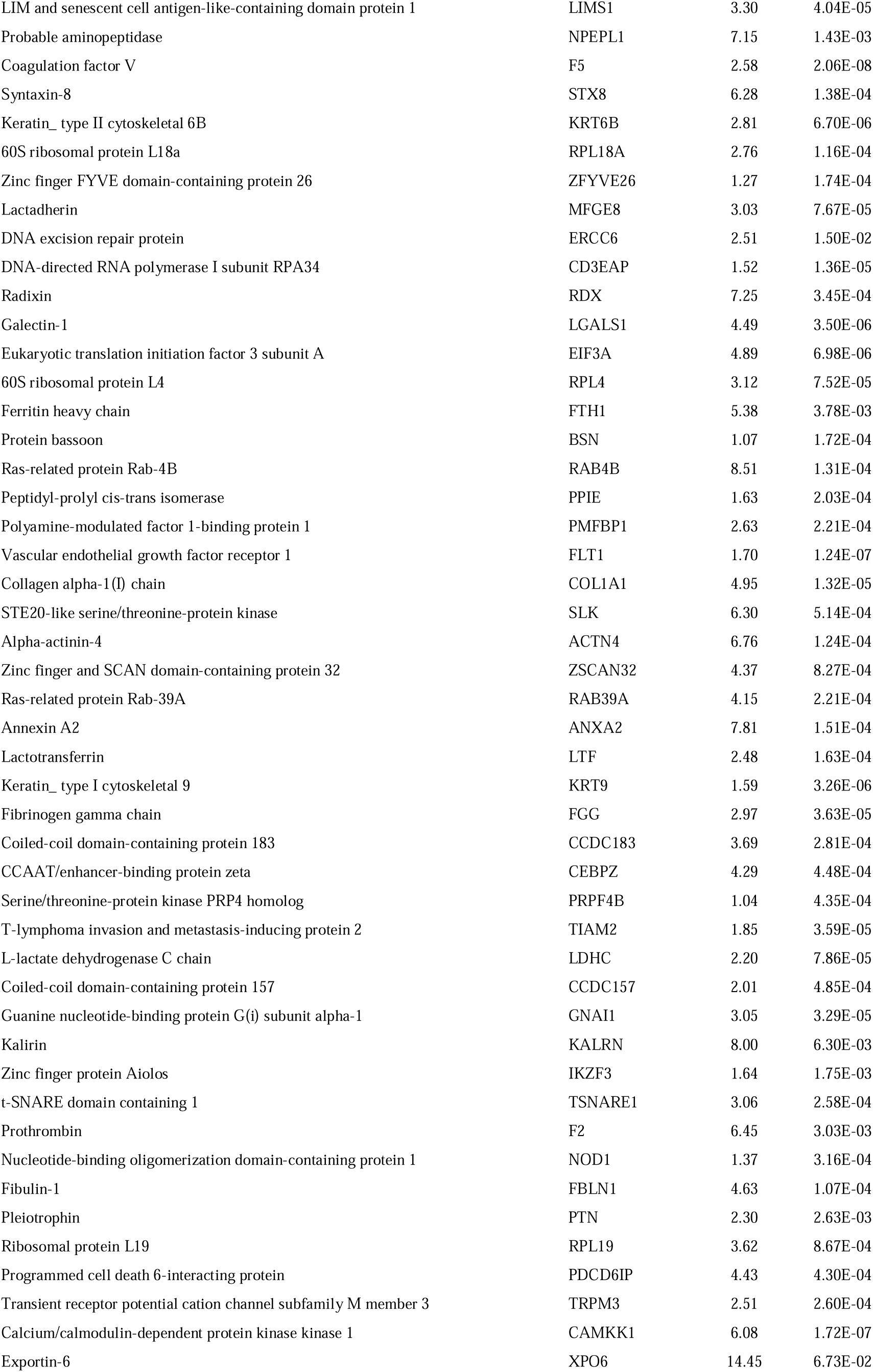
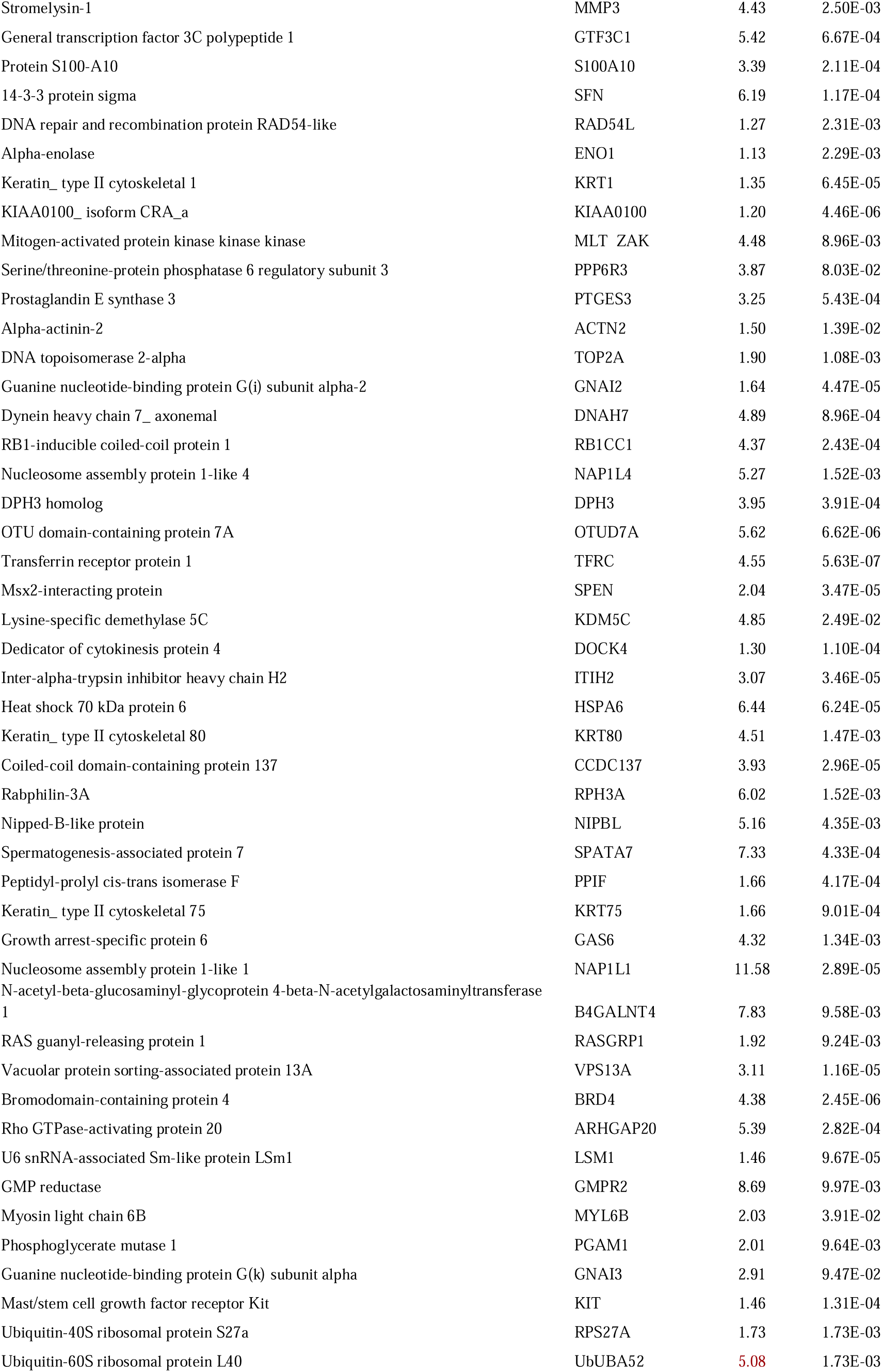
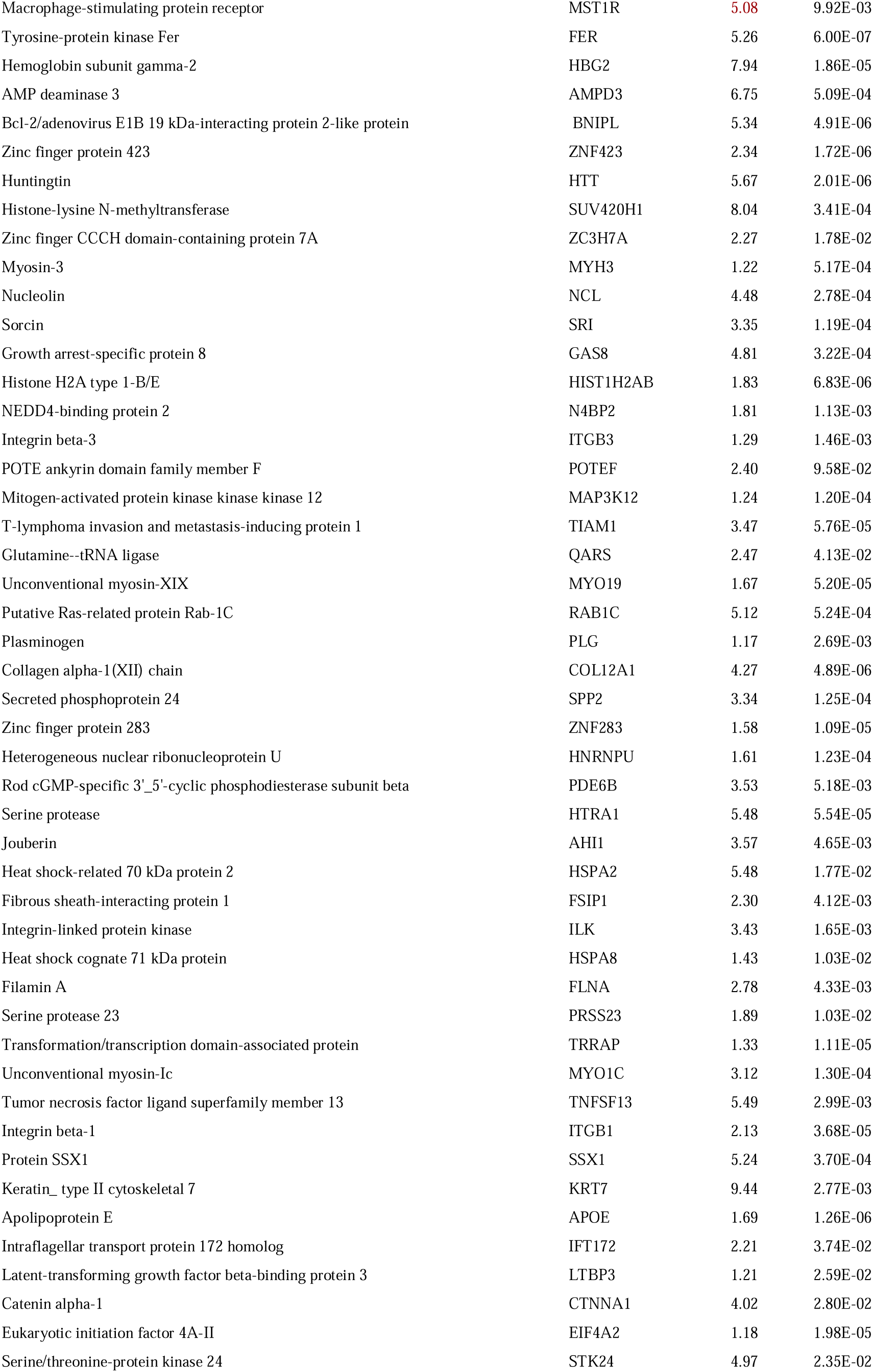
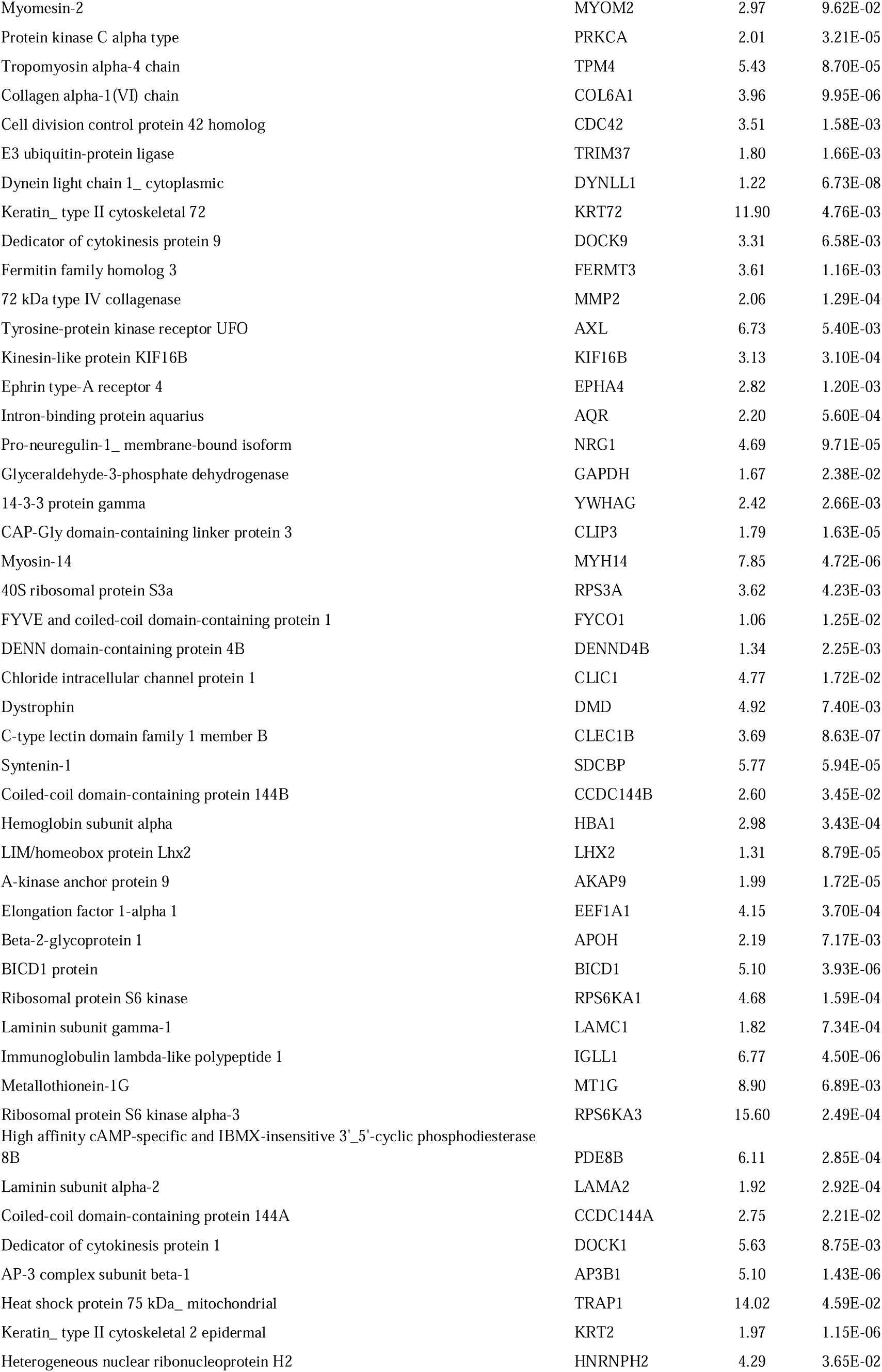
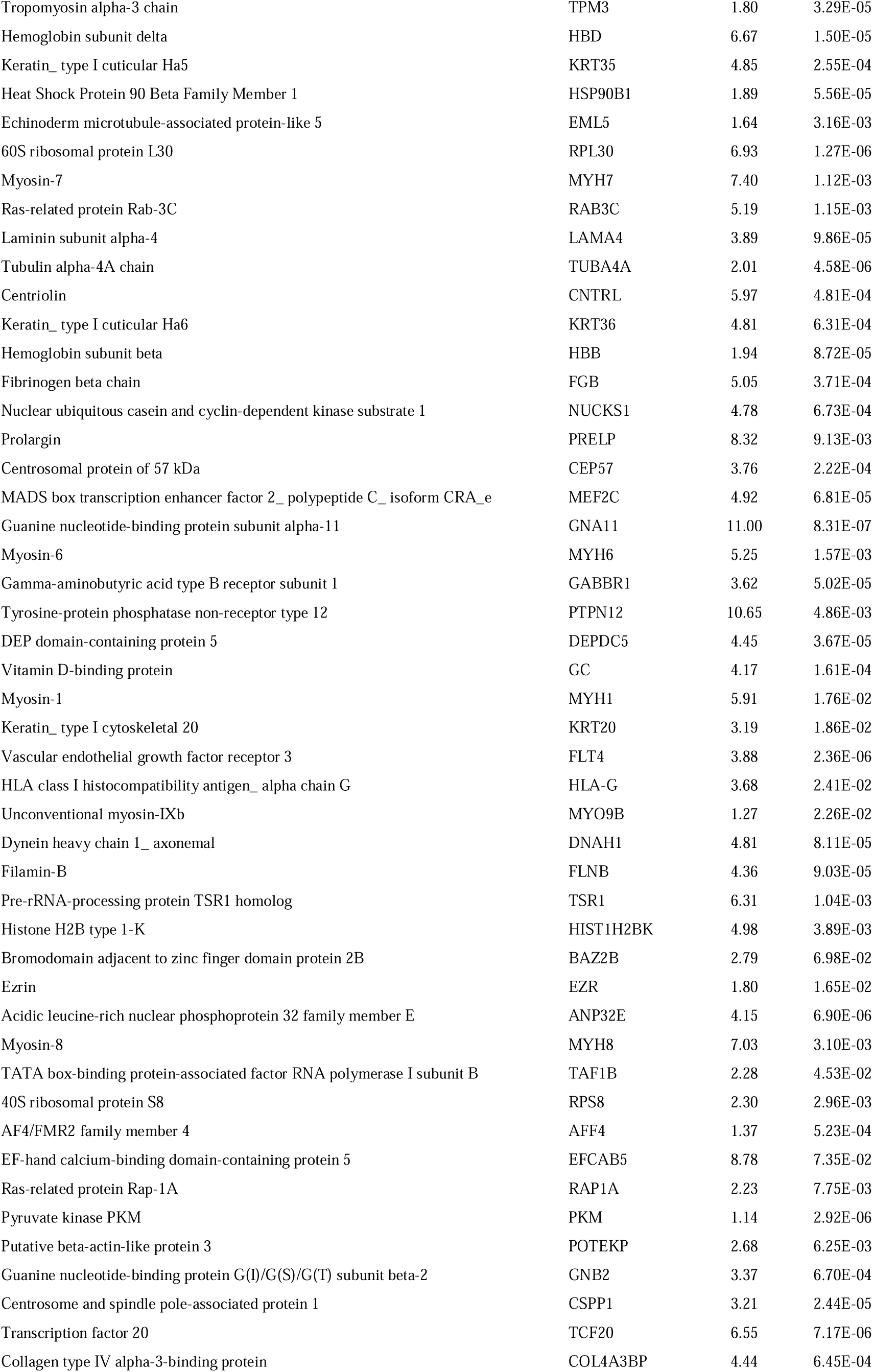
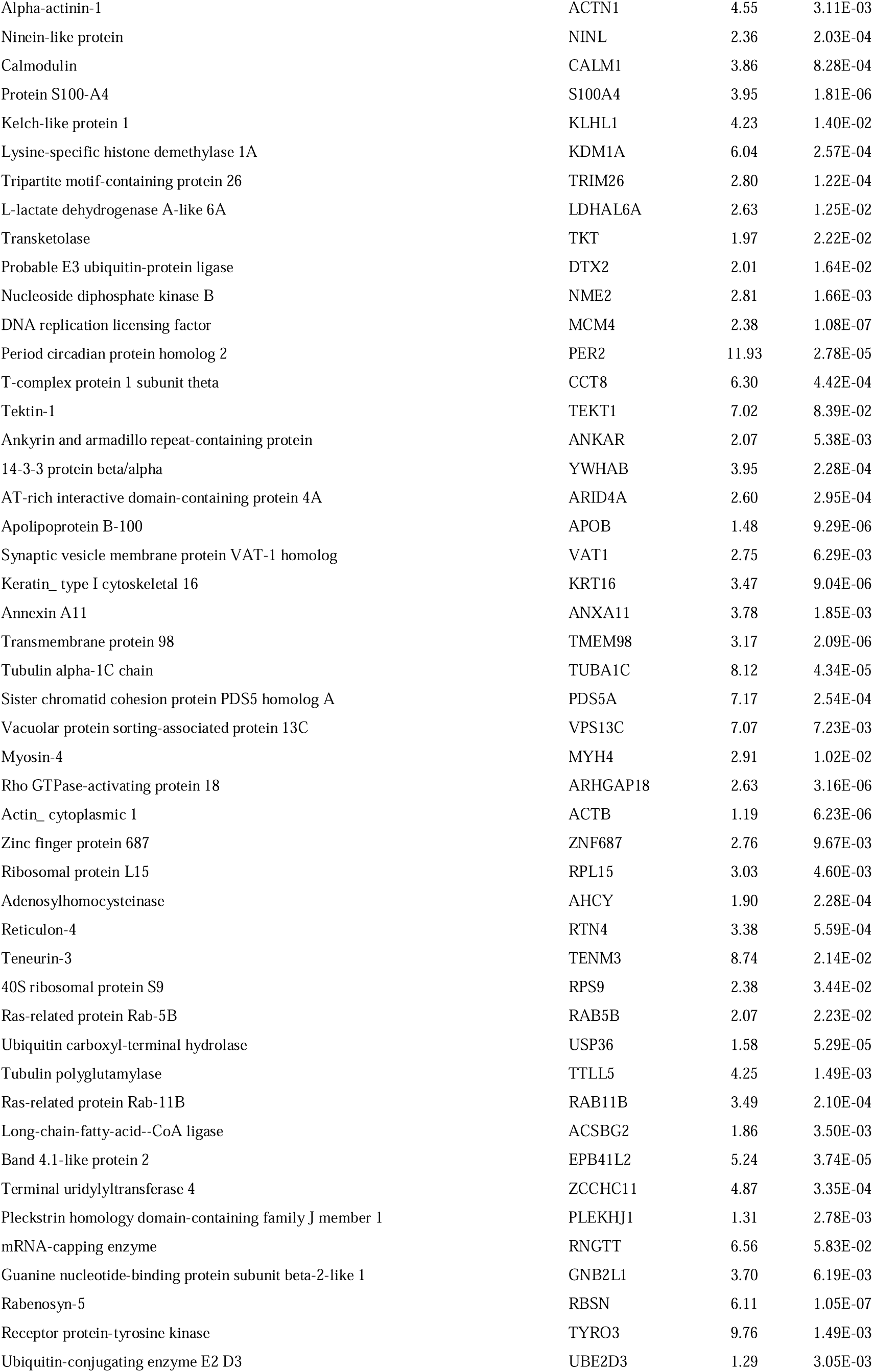
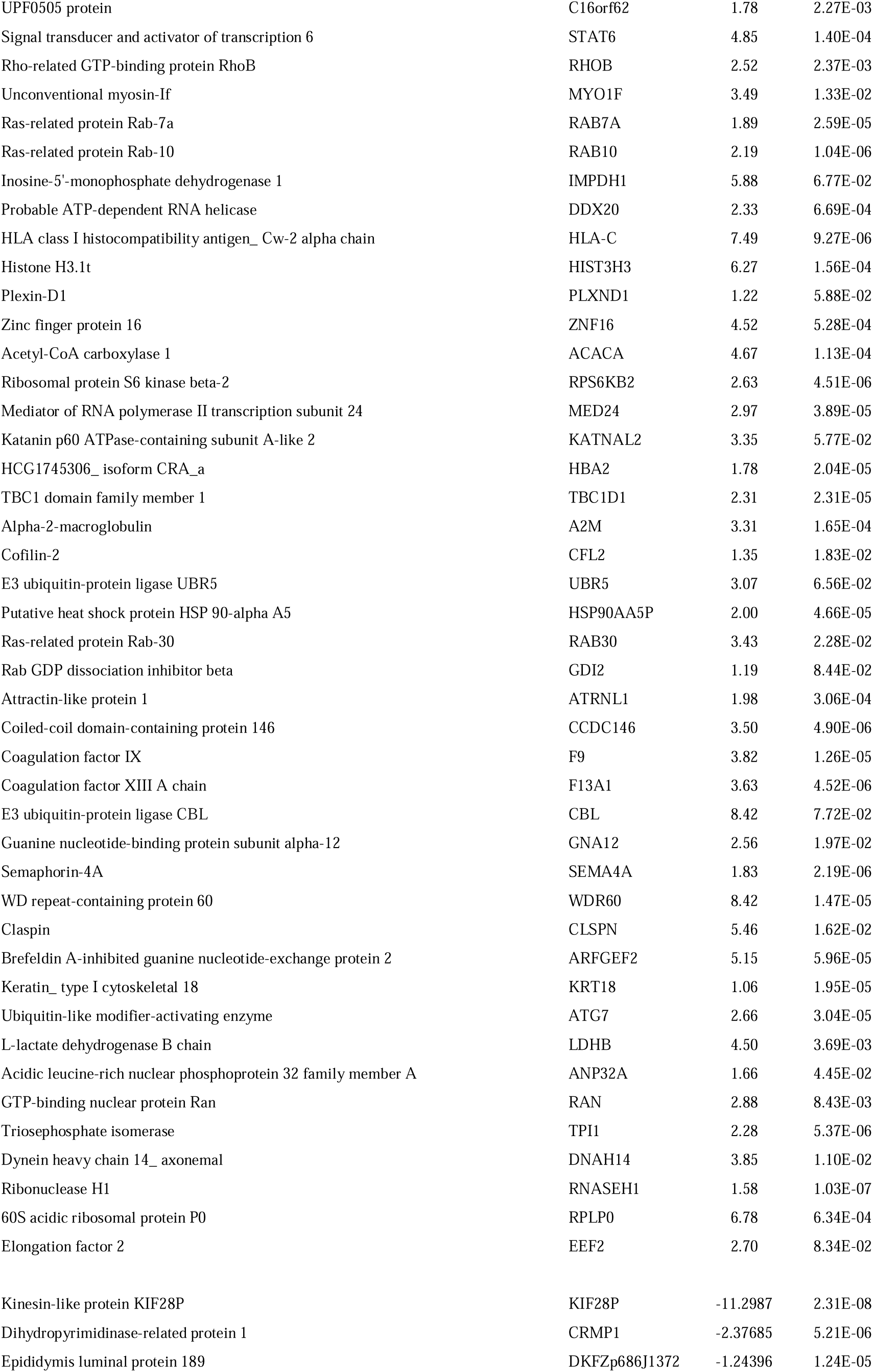
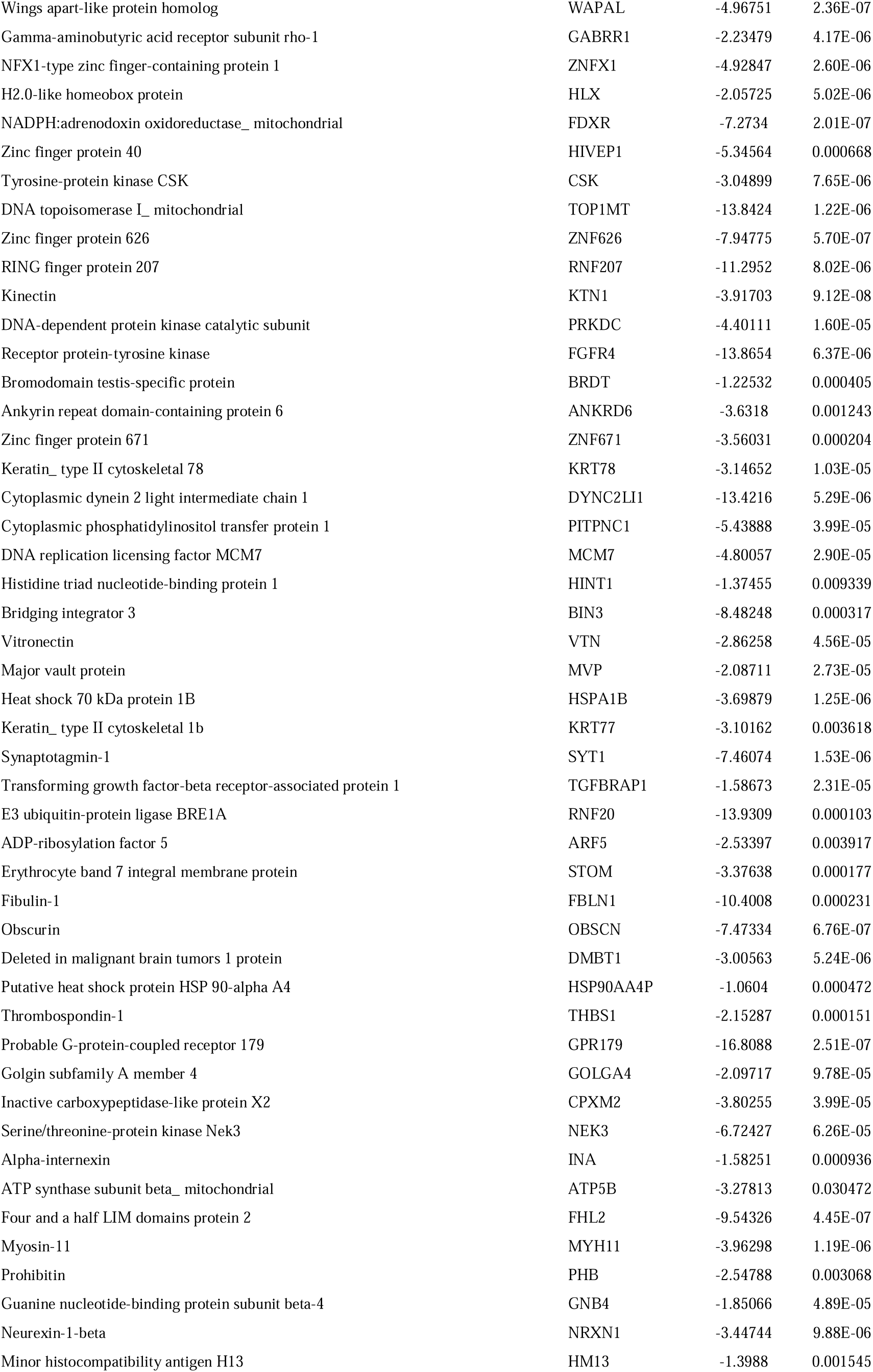
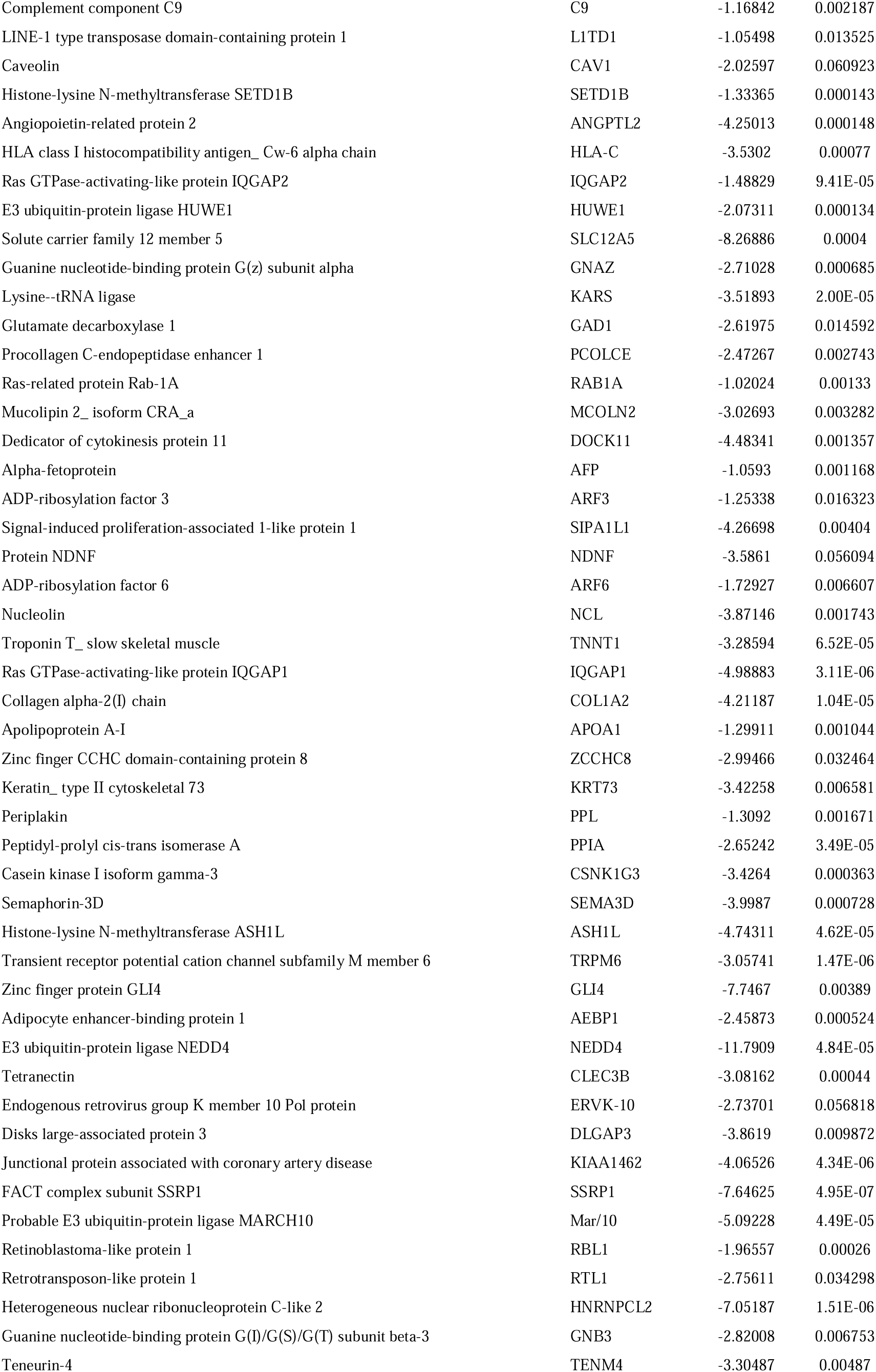
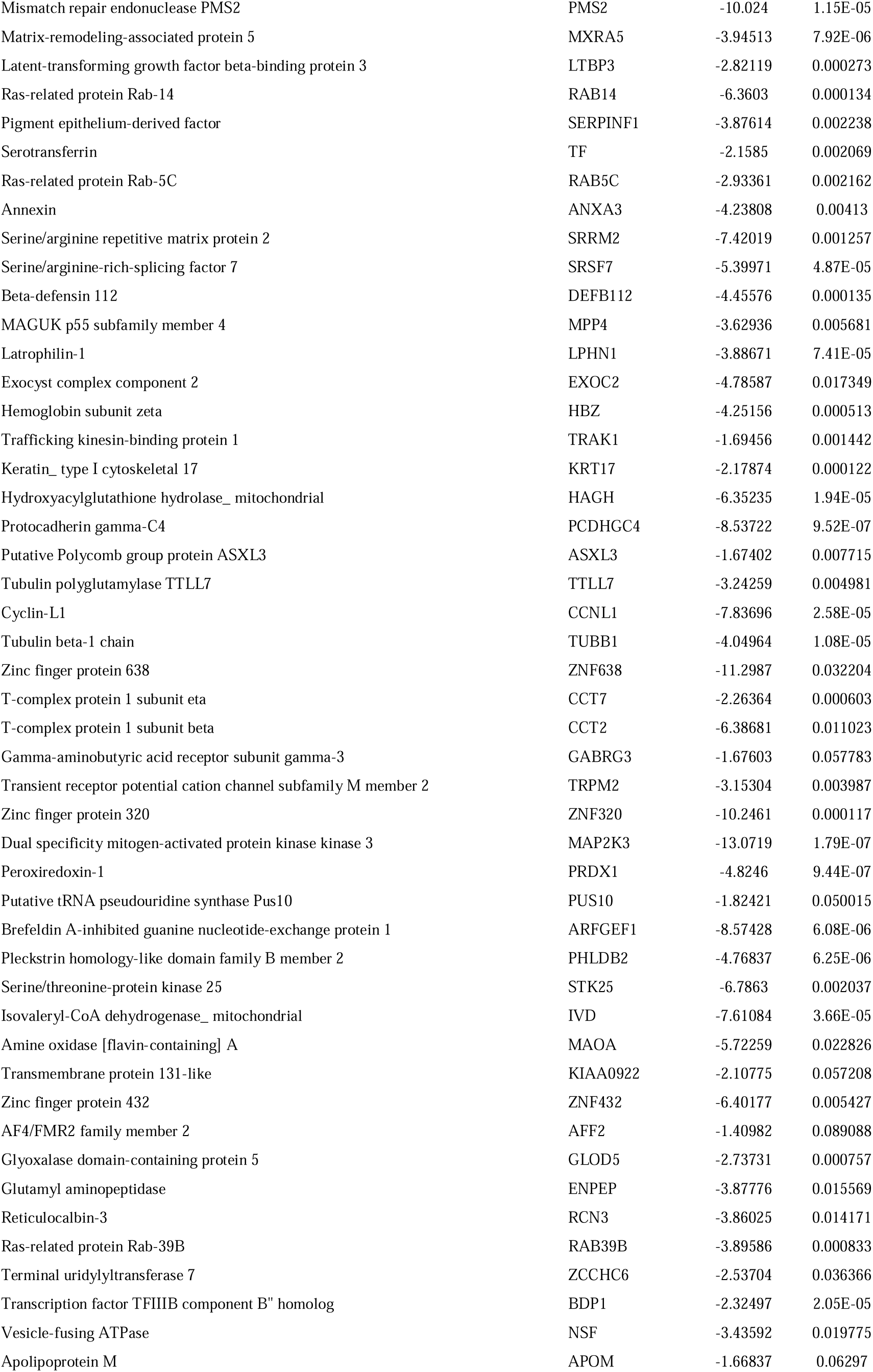
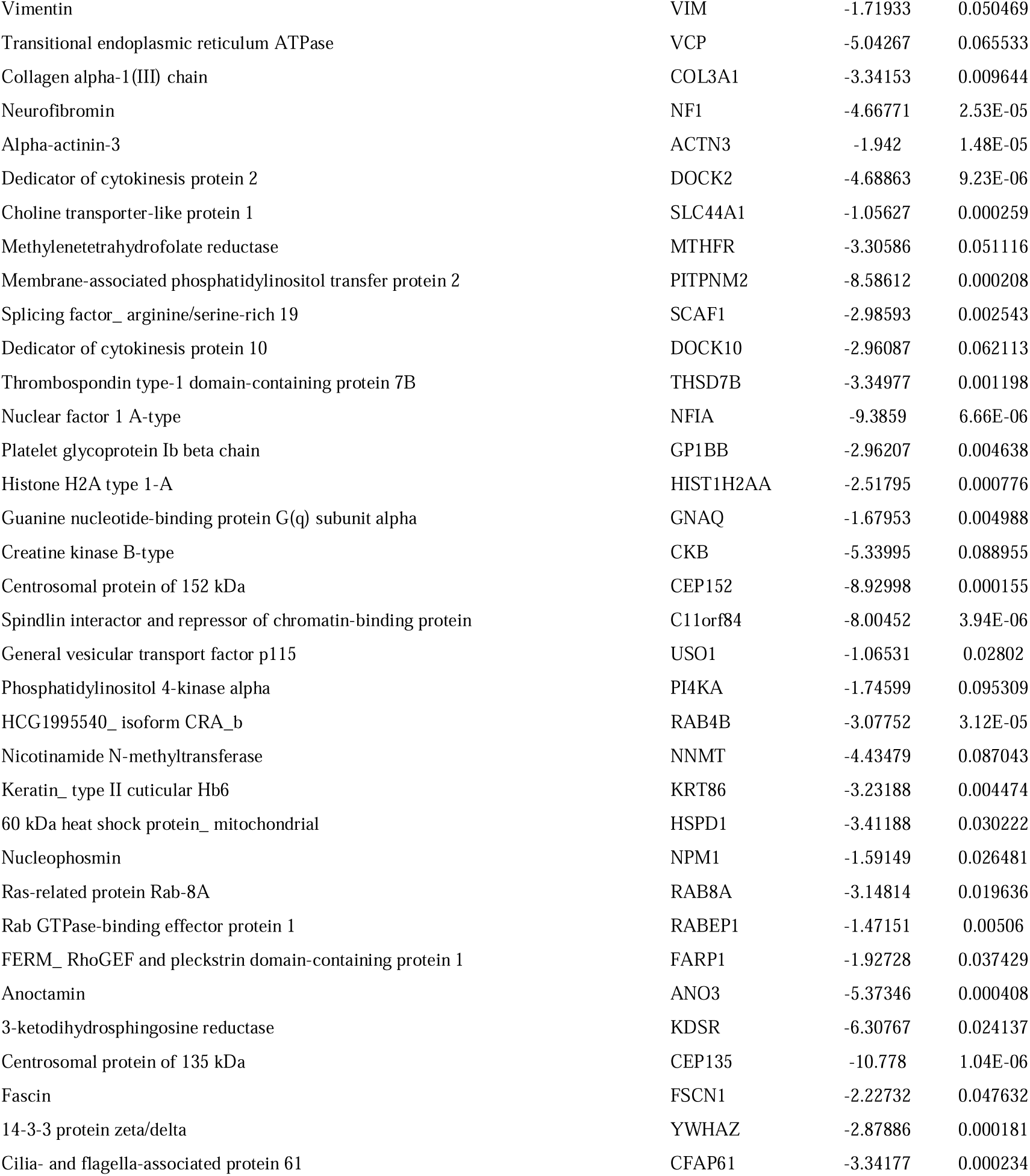
Differentially enriched proteins within the MICA/TREK EVs.

